# Yeast Rpd3L histone deacetylase couples nutrient shifts to genome-wide chromatin reprogramming

**DOI:** 10.64898/2026.02.19.706824

**Authors:** Saikat Bhattacharya, Benjamin M. Sutter, Benjamin P. Tu

**Author notes:** These authors contributed equally (Saikat Bhattacharya, Benjamin M. Sutter). Senior author (Benjamin P. Tu).

## Abstract

Adaptive transcriptional rewiring underlies the metabolic flexibility of *Saccharomyces cerevisiae*. We demonstrate that the histone deacetylase Rpd3 mediates nutrient-dependent chromatin reprogramming that coordinates transcriptional shutdown and global acetylation balance during metabolic transitions. Genome-wide analyses reveal that Rpd3 complexes drive rapid, reversible histone deacetylation across promoters and gene bodies, fine-tuning transcriptional output. Rpd3, primarily through the large complex (Rpd3L), localizes at promoters of active genes enriched in H3K9ac and the acetyltransferase Gcn5. Upon nutrient shift, Gcn5 disengages while Rpd3-mediated H3K9 deacetylation enforces repression. Loss of Rpd3 or its Rpd3L-specific subunit, Pho23, disrupts this balance, resulting in the aberrant persistence of growth programs upon starvation and defective activation of respiratory genes in the presence of glucose. HDACs thus can act as metabolic gatekeepers, coupling nutrient cues to chromatin reprogramming and ensuring transcriptional fidelity during metabolic transitions, thereby resolving the long-standing paradox of HDAC enrichment at active promoters.

## INTRODUCTION

Microorganisms exhibit remarkable plasticity in adapting to changing environments by reprogramming their metabolism and gene expression to optimize survival and growth^1,2^. In the budding yeast *Saccharomyces cerevisiae*, glucose serves as the preferred carbon source and drives rapid fermentative growth^3,4^. When glucose becomes scarce, yeast cells switch to aerobic respiration and slower growth, accompanied by extensive transcriptional rewiring^5,6^. This switch involves the coordinated regulation of hundreds of genes, enabling the repression of growth-promoting programs such as ribosome biogenesis and the activation of stress- and respiration-related pathways^7^.

Transcriptional control during nutrient transitions depends on both transcription factors and chromatin modifications. Activators such as the HAP complex promote respiratory gene expression, while repressors, including Mig1 and Mig2, inhibit genes involved in alternative carbon utilization under glucose-rich conditions^8,9^. In parallel, the chromatin landscape undergoes dynamic remodeling through histone modifications that regulate the accessibility of transcription factors and the recruitment of RNA polymerase II^10,11^. Among these modifications, histone acetylation plays a central role by modulating chromatin openness and transcriptional competence^12^. Acetylation levels reflect the balance between the activities of histone acetyltransferases (HATs) and histone deacetylases (HDACs)^13–15^. Importantly, because acetylation requires acetyl-CoA—the key metabolic intermediate linking energy status to chromatin regulation—histone acetylation directly connects transcriptional activity to cellular metabolism^16–18^.

Our previous work demonstrated that the HAT Gcn5, the catalytic subunit of the SAGA complex, redistributes the activating histone mark H3K9ac from growth-promoting genes to fatty acid oxidation (FAO) genes upon glucose depletion^19^. This redistribution is accompanied by a striking global loss of histone acetylation, suggesting that HDACs actively remove acetyl groups from chromatin during nutrient transitions. However, the mechanisms and biological significance of this widespread deacetylation, as well as the coordination of HDACs with HATs, remain unclear.

The HDAC Rpd3, a conserved class I enzyme, plays a central role in transcriptional repression and chromatin homeostasis. Intriguingly, genome-wide analyses have repeatedly shown that Rpd3 and other HDACs are enriched at actively transcribed genes^20–22^— a paradox that challenges the classical view of HDACs as mere silencers. Rpd3 forms at least two major complexes that are believed to have distinct genomic targets and functions: Rpd3L (large) and Rpd3S (small)^23,24^. Rpd3L primarily localizes to promoters, where it mediates gene repression and transcriptional memory, while Rpd3S targets gene bodies to prevent cryptic transcription^25,26^. Yet, how these complexes integrate metabolic cues to coordinate local and global acetylation dynamics remains unknown.

Here, we show that Rpd3L functions as a promoter-poised HDAC that enforces transcriptional repression during carbon source transitions. Mechanistically, Rpd3L is positioned at the promoters of active genes engaged by Gcn5, where it mediates the deacetylation of H3K9 and repression of transcription in response to changes in nutrient conditions. Loss of Rpd3 or its Rpd3L-specific subunit Pho23 disrupts this regulatory balance, resulting in aberrant gene expression and impaired metabolic adaptation. Rpd3S, on the other hand, is largely responsible for bulk histone deacetylation. Our data uncovers how an HDAC couples metabolism to chromatin regulation, ensuring appropriate transcriptional responses during nutrient fluctuations and redefining its role from a static repressor to a dynamic regulator of transcriptional output.

## RESULTS

### Rpd3 mediates nutrient-dependent histone deacetylation

To assess how perturbations in chromatin acetylation influence the metabolic adaptability of cells, we first analyzed the transcriptome of *S. cerevisiae* growing in glucose-replete conditions (+D) and following glucose depletion (–D). In glucose-rich media, transcripts associated with ribosome biogenesis and amino acid biosynthesis were among the most highly expressed, consistent with active growth and fermentative metabolism [Supplementary figure S1a]. Conversely, pathways belonging to the peroxisome that are required for fatty acid oxidation (FAO) and the utilization of non-glucose carbon sources exhibited low expression levels [Supplementary figure 1a]. Upon glucose depletion, this transcriptional profile was inverted: expression of cell-cycle and ribosomal genes sharply declined, while genes involved in respiration, gluconeogenesis, and oxidative metabolism were strongly induced [Supplementary figure S1b-d]. These observations confirm that yeast transcriptional programs mirror metabolic demand, enabling precise adaptation to nutrient availability.

To evaluate how histone modifications reflect these transcriptional transitions, we analyzed the distribution of the activating histone mark H3K9ac under both nutrient conditions. Consistent with its role in transcriptional activation, H3K9ac was enriched at the actively transcribed genes discussed above in both +D and –D states [Figure 1a]. Moreover, in both conditions, the extent of H3K9ac enrichment correlated positively with gene expression levels, confirming its tight association with transcriptional activity [Figure 1b]. These findings demonstrate that histone acetylation undergoes redistribution in response to nutrient shifts while maintaining strong alignment with active transcriptional programs.

**Figure 1:**
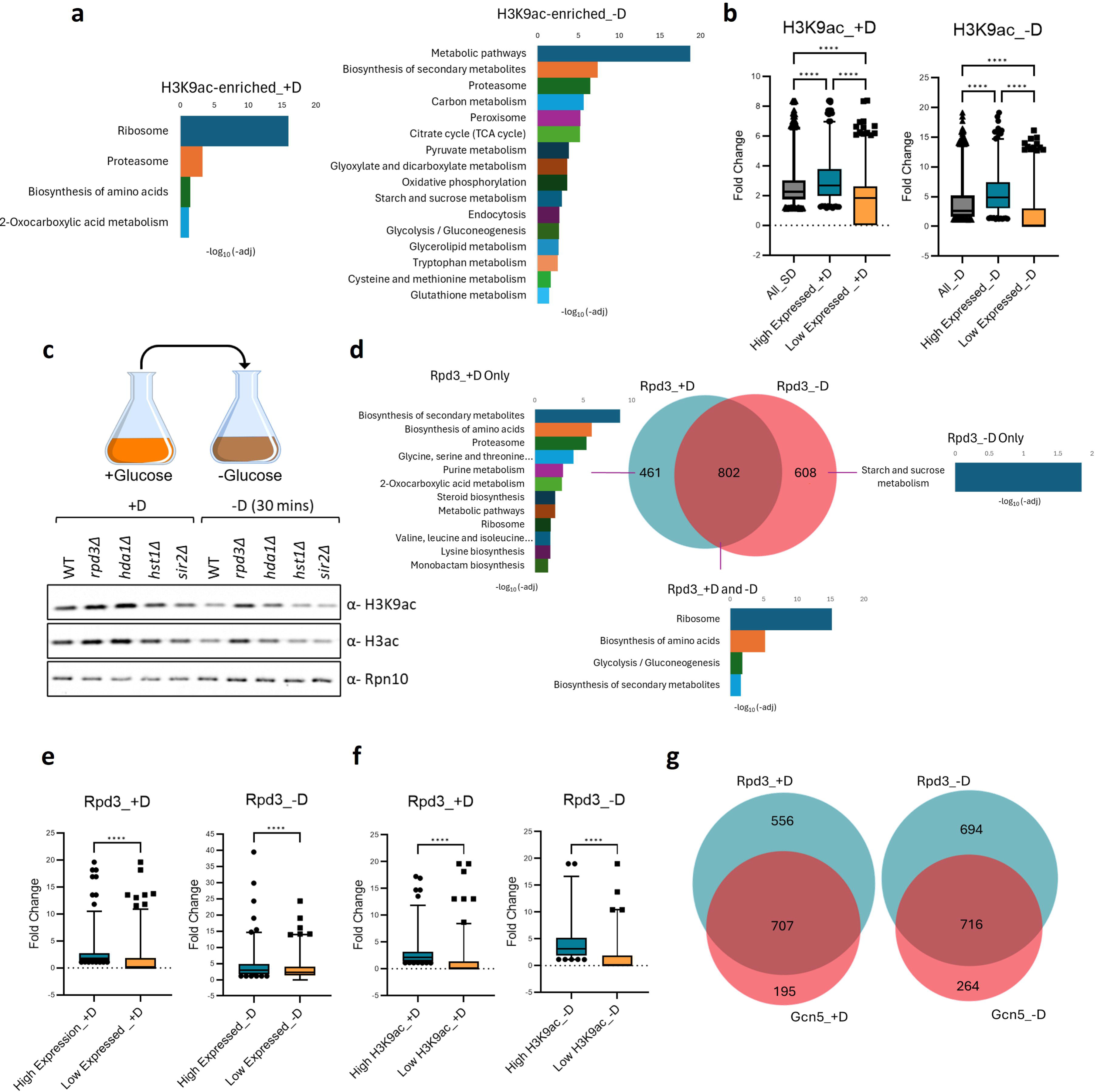
Rpd3 associates with transcriptionally active genes under nutrient-dependent conditions. (a) Chromatin immunoprecipitation followed by sequencing (ChIP-seq) was performed to profile genome-wide enrichment of the histone H3 lysine 9 acetylation (H3K9ac) mark in *Saccharomyces cerevisiae* (CEN.PK strain) grown in glucose-replete (+D) or glucose-depleted (−D) synthetic media. Genes enriched for H3K9ac under each condition were subjected to KEGG pathway analysis. In both nutrient conditions, H3K9ac is preferentially enriched at genes that are transcriptionally active, with distinct pathway enrichment reflecting metabolic adaptation to glucose availability. (b) Box plots showing the distribution of H3K9ac enrichment (expressed as fold change over background) across genes stratified by expression level (high vs. low) in +D and −D conditions. In both media conditions, highly expressed genes exhibit significantly greater H3K9ac enrichment compared to lowly expressed genes, supporting a strong correlation between H3K9 acetylation and transcriptional activity. Ordinary one-way ANOVA was used to calculate statistical significance. ****p<0.0001. (c) Wild-type (WT) and mutant yeast strains were grown in glucose-replete (+D) media to mid-exponential phase (OD600 ≈ 1), followed by a shift to glucose-depleted (−D) media for 30 minutes. Whole-cell lysates of yeast were prepared and analyzed by western blot to assess global levels of H3K9ac and total H3 acetylation (H3ac). Rpn10 was probed as the loading control. Deletion of *RPD3* resulted in the highest retention of H3K9ac levels upon glucose depletion, indicating that Rpd3 contributes to histone deacetylation during metabolic stress. (d) ChIP-seq analysis of Rpd3 occupancy was performed in cells grown under +D and −D conditions. A Venn diagram illustrates the overlap and condition-specific binding of Rpd3 target genes. Genes unique to each condition, as well as shared targets, were subjected to KEGG pathway analysis, and enriched pathways are indicated. Rpd3 exhibits dynamic, condition-dependent chromatin association, preferentially binding to distinct subsets of genes that are transcriptionally active under each metabolic state. (e, f) Box plots showing Rpd3 enrichment (fold change over background) across genes categorized based on (e) transcriptional output (high vs. low expression) and (f) H3K9ac enrichment levels. Ordinary one-way ANOVA was used to calculate statistical significance. ****p<0.0001. In both analyses, Rpd3 shows higher occupancy at genes with elevated transcription and increased H3K9ac levels, suggesting a role for Rpd3 in modulating chromatin at actively transcribed loci. (g) Venn diagram derived from ChIP-seq datasets comparing genomic targets of Rpd3 and the histone acetyltransferase Gcn5 under +D and −D conditions. The overlap highlights both shared and distinct target genes, providing insight into the coordinated and potentially antagonistic roles of these chromatin modifiers in regulating transcription in response to nutrient availability.

We previously demonstrated that histone H3 undergoes rapid deacetylation upon shifting cells from +D to –D^19^. Yeast contains multiple HDACs belonging to three classes [Supplementary figure S2a]. To test whether Rpd3 is primarily responsible for the global deacetylation observed upon starvation, we generated deletion strains for multiple HDACs spanning different functional classes. We monitored histone acetylation levels during the +D to –D transition. Among the tested mutants, the loss of Rpd3 had the most prominent effect on the starvation-induced decrease in not only H3K9ac but also H3ac levels in general, indicating that it is the principal deacetylase mediating this process [Figure 1c]. These results identify Rpd3 as a key effector of global chromatin deacetylation in response to glucose deprivation.

### Rpd3 binding correlates with transcriptionally active genes and Gcn5 occupancy

Glucose deprivation triggers a rapid, genome-wide deacetylation of histones mediated by Rpd3, which is quickly reversed upon re-exposure to glucose^19^. To elucidate the mechanistic basis of this response, we performed Rpd3 chromatin immunoprecipitation followed by sequencing (ChIP-seq) under both nutrient conditions. Rpd3 displayed dynamic binding behavior, associating with distinct gene sets in +D and –D, yet approximately two-thirds of Rpd3-bound loci were shared between the two states [Figure 1d].

Analysis of genes that are Rpd3-bound only in +D or –D revealed that, in both nutrient states, Rpd3 preferentially occupied highly expressed genes. In +D, Rpd3 bound to genes belonging to growth pathways such as the biosynthesis of secondary metabolites and amino acids, whereas in –D, Rpd3 was enriched at genes required for the utilization of alternate carbon sources [Figure 1d]. Moreover, a significant positive correlation was observed between Rpd3 binding and transcript abundance. Rpd3 showed higher enrichment at highly expressed genes than at lowly expressed ones, underscoring Rpd3’s unexpected association with transcriptionally active loci [Figure 1e].

To further examine this relationship, we compared Rpd3’s enrichment with that of the activating H3K9ac mark. Rpd3 binding showed a preference for regions with elevated H3K9ac, a pattern consistent across both nutrient states [Figure 1f]. Given that H3K9ac is deposited by the histone acetyltransferase Gcn5, we next assessed the degree of overlap between Rpd3 and Gcn5 occupancy. Genome-wide co-localization analyses revealed a substantial overlap between the two, suggesting that Rpd3 and Gcn5 co-occupied genes [Figure 1g]. Moreover, the co-occupied genes were largely enriched in growth and respiratory pathways in +D and -D, respectively [Supplementary figure S2b].

Despite the overlap in the binding between Rpd3 and Gcn5, some clear differences also emerged upon analyzing the genes bound by Gcn5. Gcn5 binds distinct but overlapping gene sets in +D and –D, with enrichment at highly active genes—consistent with its role in depositing H3K9ac [Supplementary figure S2c]. Interestingly, while both enzymes target active genes, their persistent binding sites differed: Rpd3 remained associated with ribosomal genes across conditions [Figure 1d], whereas Gcn5 only bound to them in +D [Supplementary figure S2c]. Also, Gcn5 showed sustained enrichment at metabolic gene promoters [Supplementary figure S2c]. Thus, while Rpd3 and Gcn5 share overlapping occupancy patterns, their binding dynamics diverge in gene-category specificity, hinting at opposing yet coordinated regulatory functions.

### Loss of Rpd3 disrupts transcriptional rewiring and induces metabolic misexpression

Although Rpd3 is classically regarded as a transcriptional repressor, its enrichment at active promoters prompted us to test whether its regulatory outcome mirrors that of Gcn5, a transcriptional activator. We therefore compared the transcriptomes of *rpd3Δ* and *gcn5Δ* strains under glucose-replete conditions (+D). Notably, the overlap between genes up- or downregulated in each mutant was minimal [Supplementary figure S3a, b]. Also, only six genes were common among the downregulated set in *gcn5Δ* and the upregulated set in *rpd3Δ* out of more than 400 total, indicating that despite their co-occupancy, the two enzymes have distinct effects on transcriptional outcomes [Figure 2a].

**Figure 2:**
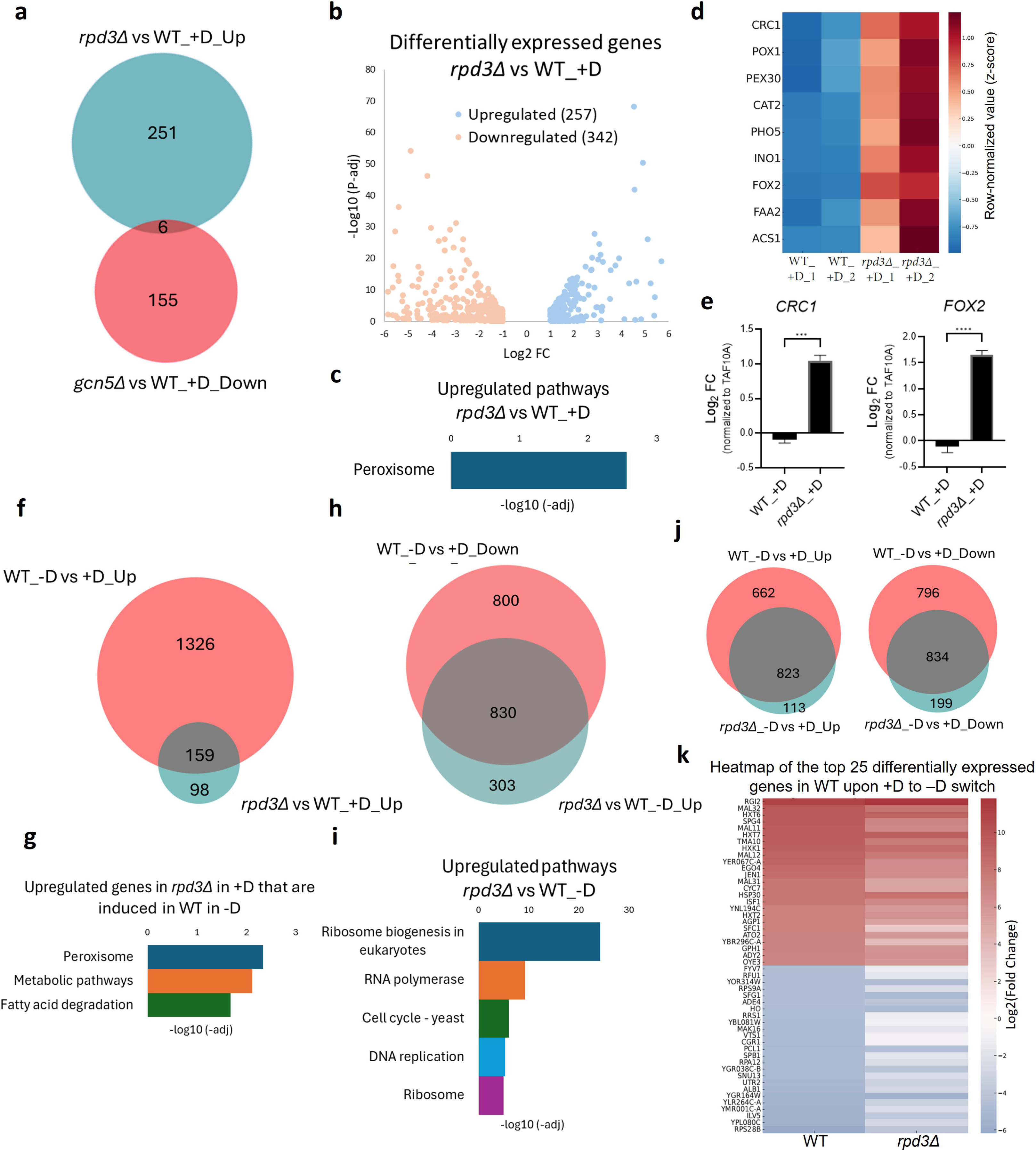
Rpd3 is required for transcriptional rewiring upon metabolic transitions. (a) Venn diagram illustrating the overlap between genes that are significantly upregulated in *rpd3Δ* cells grown in glucose-replete conditions (+D; fold change ≥2, FDR <0.05) and genes that are downregulated in *gcn5Δ* cells under the same condition. The meagre overlap highlights the antagonistic roles in transcriptional regulation despite a significant overlap in chromatin binding. (b) Volcano plot depicting genome-wide differential gene expression in *rpd3Δ* versus WT cells grown in +D. Each dot represents a gene, plotted by log2 fold change versus −log10(FDR). Significantly upregulated and downregulated genes are highlighted, illustrating widespread transcriptional perturbation upon loss of Rpd3. (c) KEGG pathway enrichment analysis of genes upregulated in *rpd3Δ* compared to WT in +D conditions. The enriched pathway belongs to peroxisomal β oxidation, indicating that Rpd3 suppresses specific metabolic gene programs under glucose-replete conditions. (d) Heatmap showing normalized expression levels of selected genes involved in fatty acid oxidation (FAO) in WT and *rpd3Δ* cells grown in +D. FAO-related genes exhibit increased expression in *rpd3Δ*, suggesting derepression of alternative metabolic pathways in the absence of Rpd3. (e) Quantitative PCR (qPCR) validation of representative FAO genes, *CRC1* and *FOX2*, in WT and *rpd3Δ* cells under +D conditions. Gene expression levels were normalized to the housekeeping gene *TAF10A*. Ordinary one-way ANOVA was used to calculate statistical significance. n=3, ***p=0.0002, ****p<0.0001. The error bars depict the Standard Error of Mean (SEM). Consistent with RNA-seq results, both *CRC1* and *FOX2* show elevated expression in *rpd3Δ* cells. (f, h, j) Venn diagrams comparing differentially expressed genes (DEGs) between WT and *rpd3Δ* strains across the indicated nutrient conditions (e.g., +D, −D, and glucose shift). These analyses reveal both shared and condition-specific transcriptional responses, highlighting the role of Rpd3 in coordinating gene expression changes during metabolic transitions. (g, i) KEGG pathway enrichment analyses of genes upregulated in *rpd3Δ* relative to WT under the corresponding conditions shown in (f, h, j). Enriched pathways reflect metabolic and stress-responsive processes that are aberrantly activated in the absence of Rpd3. (k) Heatmap displaying the top 25 most upregulated and top 25 most downregulated genes in WT cells upon transition from +D to glucose-depleted −D conditions. The expression patterns of these genes are shown for both WT and *rpd3Δ* strains, illustrating that loss of Rpd3 impairs proper transcriptional reprogramming during nutrient shifts.

In *gcn5Δ* cells, genes that were normally highly expressed in +D—primarily ribosomal and growth-related genes—were significantly downregulated, consistent with the activator role of Gcn5 [Supplementary figure S3c]. Upon glucose depletion (–D), *gcn5Δ* cells failed to induce key metabolic and respiratory genes, confirming Gcn5’s role in enabling transcriptional activation during nutrient transitions [Supplementary figure S3d].

In contrast, Rpd3 deletion affected 599 genes in glucose, belonging to various metabolic pathways, with 257 showing derepression [Figure 2b, supplementary figure S3e]. This includes the known Rpd3 target *INO1* as well as genes such as *SPO13*, *IME1*, *CAR2,* and *PHO5,* which have been previously reported to be upregulated upon Rpd3 deletion^27^ [Table S1]. These derepressed genes were enriched for peroxisomal pathways and included genes such as *CRC1*, *CAT2,* and *FOX2* that result in fatty acid oxidation—processes normally suppressed in glucose-rich conditions but induced upon starvation [Figure 2c, d]. qPCR validation confirmed the derepression of these genes in *rpd3Δ* cells grown in glucose, supporting Rpd3’s role in their repression [Figure 2e]. Notably, more than 60% of genes derepressed in *rpd3Δ* overlapped with those that are normally induced in wild-type (WT) cells upon glucose depletion [Figure 2f]. These common genes also represented the peroxisome and fatty acid degradation pathways, emphasizing that Rpd3 suppresses metabolic programs in +D that should only be activated during glucose scarcity (-D) [Figure 2g].

Upon shifting to –D, *rpd3Δ* cells exhibited extensive misregulation, with 1,898 genes differentially expressed—1,133 upregulated and 765 downregulated [Supplementary figure S3f]. A closer inspection of these genes revealed the same theme observed in +D—aberrant expression of metabolic programs. Over 70% of genes upregulated in *rpd3Δ* in –D corresponded to those normally downregulated in WT cells upon glucose withdrawal [Figure 2h]. The upregulated genes in *rpd3Δ* in –D included ribosomal, cell-cycle, and DNA replication pathways, which are normally repressed in WT cells under starvation [Figure 2i]. Moreover, the downregulated genes belonged to pathways such as oxidative phosphorylation and utilization of alternative carbon sources, suggesting that *rpd3Δ* interferes with the respiratory adaptation of cells [Supplementary figure S3g].

Indeed, comparison of +D to –D transitions revealed that while WT cells showed robust transcriptional rewiring, *rpd3Δ* cells exhibited blunted responses, with fewer genes undergoing differential expression and reduced fold-change magnitudes [Figure 2j, k]. These findings indicate that Rpd3 is essential for the transcriptional shutdown of growth programs and proper induction of metabolic genes during nutrient transitions.

### Rpd3 exhibits nutrient-dependent chromatin occupancy and distinct transcriptional regulatory signatures

Unlike Gcn5, whose deletion caused downregulation of its direct target genes, the pathways upregulated in the *rpd3Δ* strain did not align with the prominent genomic enrichment of Rpd3 binding. This discrepancy suggested that Rpd3’s regulatory role may depend on both the identity of its targets and the chromatin context in which it acts. To examine this, we profiled Rpd3’s genome-wide occupancy under glucose-rich (+D) and glucose-depleted (–D) conditions.

A metagene analysis of Rpd3’s genome-wide binding revealed that it undergoes a pronounced redistribution between nutrient states [Figure 3a, b]. Under glucose-replete conditions, strikingly, Rpd3 occupied both promoters and gene bodies, indicating broad chromatin engagement during active proliferation [Figure 3a]. In +D, close to a quarter of Rpd3-binding sites were in the gene body (exons), while three-quarters of the binding sites were at the promoters [Figure 3c]. In contrast, the activating HAT Gcn5 was mostly enriched at the promoters [Figure 3a, c]. In –D, however, more than 99% of Rpd3 binding was concentrated at promoter regions upstream of transcription start sites (TSSs), similar to the pattern of Gcn5 [Figure 3b, d].

**Figure 3:**
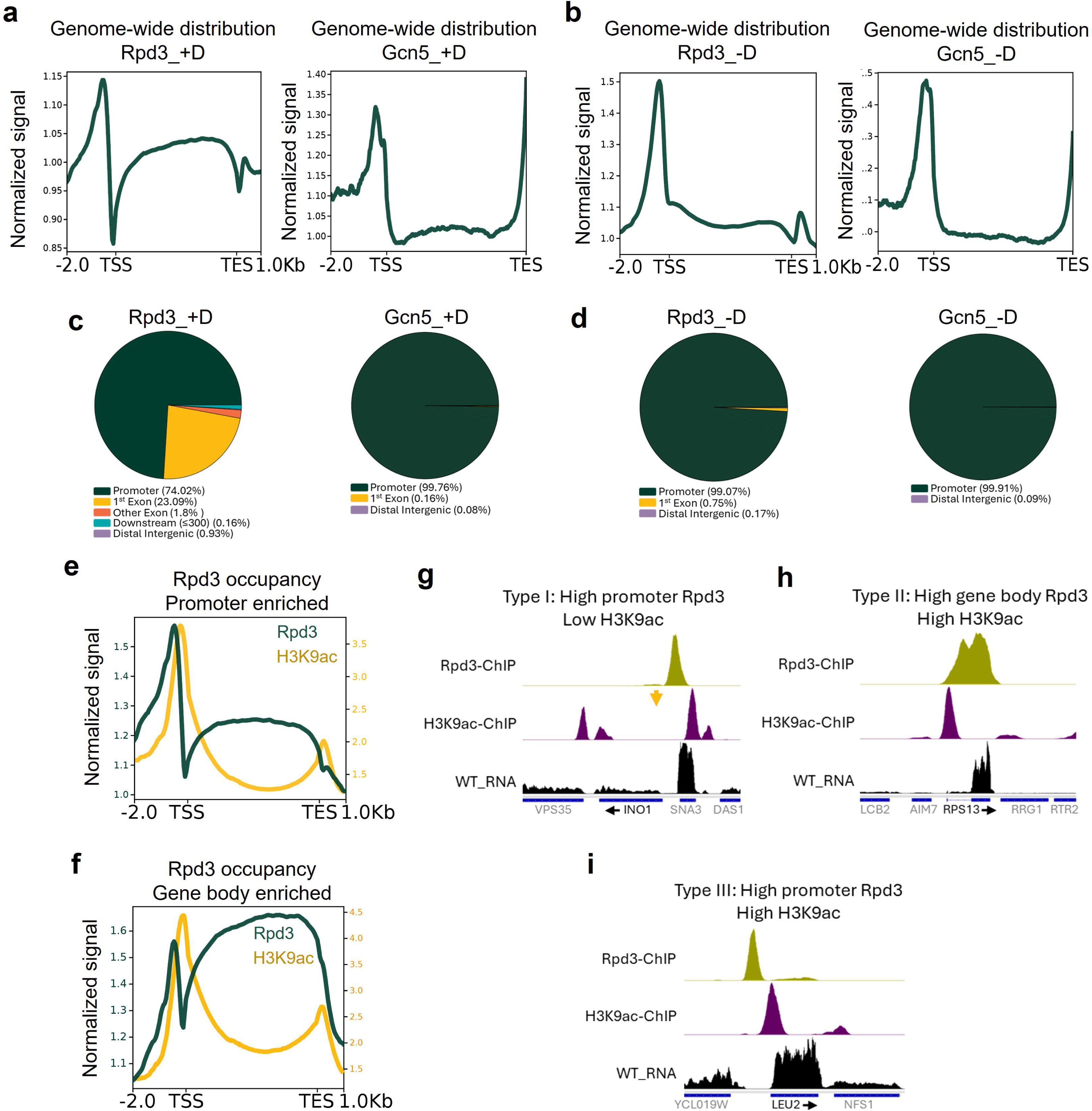
Rpd3-bound genes exhibit distinct chromatin and regulatory features. (a, b) Metagene profiles showing the genome-wide distribution of Rpd3 and Gcn5 across gene bodies and flanking regions in cells grown under glucose-replete (+D) and glucose-depleted (−D) conditions. TSS = Transcription Start Site, TES = Transcription End Site. Gcn5 displays a strong and consistent enrichment at promoter regions regardless of nutrient availability. In contrast, Rpd3 exhibits a broader distribution in +D conditions, with enrichment at both promoter regions and across gene bodies, indicating a more diverse mode of chromatin association. (c, d). Pie charts summarizing the genomic distribution of Rpd3- and Gcn5-bound regions across different gene elements (e.g., promoters, gene bodies, and intergenic regions) in +D and −D conditions. Gcn5 binding is predominantly restricted to promoter regions in both conditions. In contrast, only ∼74% of Rpd3-bound sites localize to promoters in +D, with the remaining fraction distributed across gene bodies and other genomic regions, highlighting its distinct and more widespread binding pattern. (e, f) Metagene plots showing H3K9ac enrichment at genes stratified based on Rpd3 binding location (promoter-enriched versus gene body-enriched). Regardless of whether Rpd3 is localized to promoters or gene bodies, its binding sites are associated with elevated H3K9ac levels, indicating that Rpd3 occupancy correlates with transcriptionally active chromatin states. (g-i) Representative genome browser tracks displaying ChIP-seq signal for Rpd3 and H3K9ac, along with RNA-seq read coverage (CPM-normalized), at selected loci. These examples illustrate the relationship between Rpd3 binding, histone acetylation, and transcriptional output. The arrow in (g) highlights a region where Rpd3 binding is observed in the absence of a corresponding H3K9ac peak, suggesting context-dependent differences in chromatin modification at Rpd3 target sites.

Based on its positional enrichment, Rpd3-bound genes in +D could be classified into two groups: promoter-enriched and gene body–enriched [Figure 3e, f]. Both groups exhibited strong H3K9ac signal near the TSS, suggesting that Rpd3 preferentially associates with transcriptionally active loci regardless of its binding position on gene elements [Figure 3e, f].

Next, an integrative analysis of Rpd3 ChIP–seq, H3K9ac enrichment, and RNA–seq expression was performed, which defined three distinct binding signatures. Type I genes, exemplified by the ‘canonical’ Rpd3 target *INO1*, showed Rpd3 enrichment at promoters with low H3K9ac and repressed transcription, consistent with classical HDAC-mediated silencing [Figure 3g]. Type II genes, such as ribosomal loci, displayed Rpd3 occupancy across gene bodies with strong TSS-proximal H3K9ac enrichment, suggesting a potential role in transcriptional fine-tuning or suppression of cryptic initiation [Figure 3h]. Type III genes, including *LEU2*, represented an unexpected category or ‘atypical’ binding in which Rpd3 occupied promoters of highly active genes that retained robust H3K9ac near the TSS [Figure 3i].

Together, these findings demonstrate that Rpd3’s chromatin association is dynamic and context-dependent, varying with nutrient state, gene identity, and transcriptional activity. This reveals a multifaceted role for Rpd3 in balancing activation and repression across distinct genomic contexts.

### Promoter-associated Rpd3 restricts H3K9ac enrichment and transcriptional activation

To determine whether Rpd3 directly represses genes derepressed upon its loss, we compared its chromatin occupancy with transcriptional changes in the *rpd3Δ* strain. More than 22% of genes upregulated in both glucose-replete (+D) and glucose-depleted (–D) conditions exhibited Rpd3 binding, predominantly at promoter regions—consistent with the Type I, canonical regulatory mode [Figure 4a, b].

**Figure 4:**
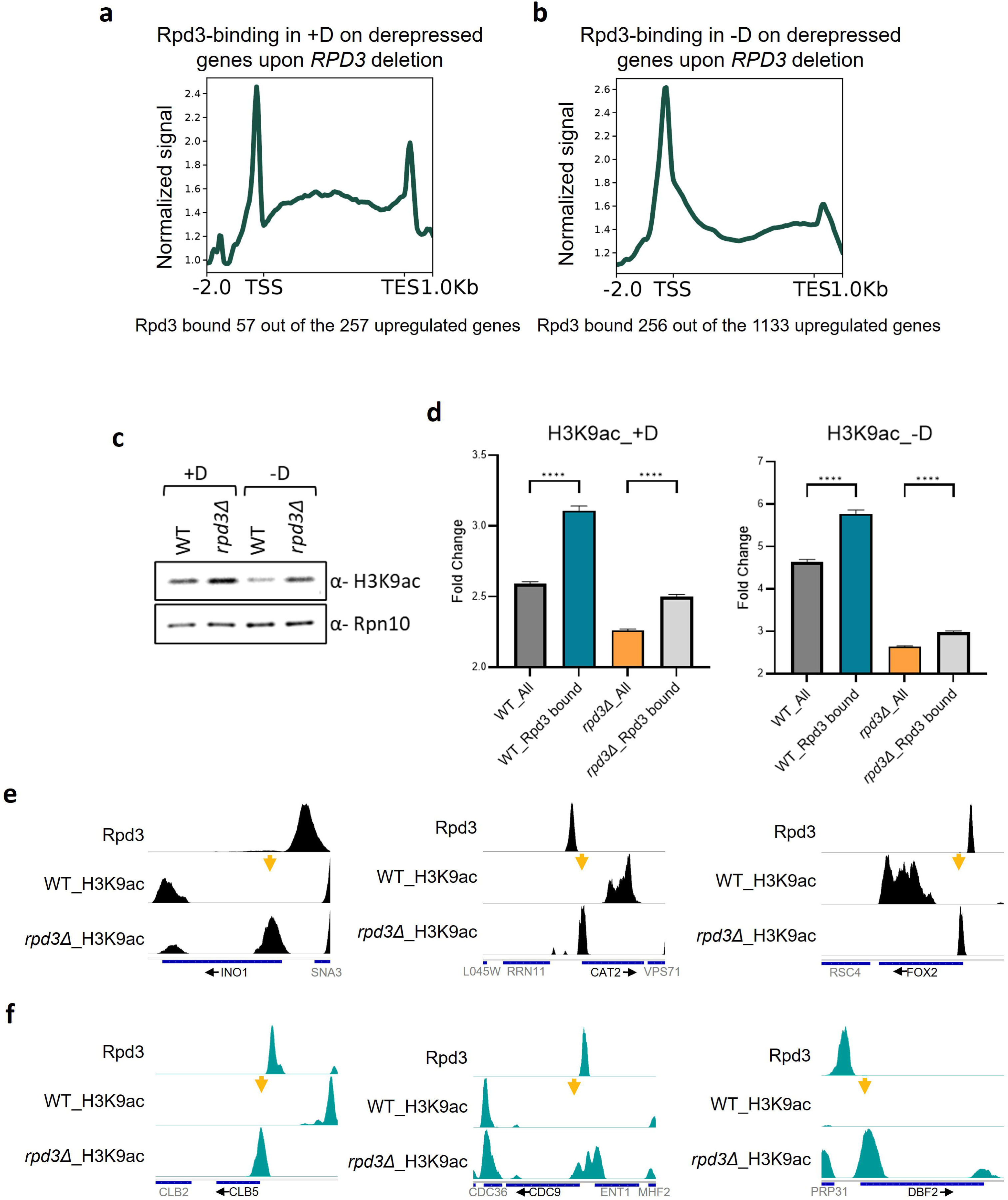
Rpd3-mediated histone deacetylation drives transcriptional repression during nutrient transitions. (a, b) Metagene profiles showing the genome-wide enrichment of Rpd3 at genes that become derepressed upon *RPD3* deletion in glucose-replete (+D) and glucose-depleted (−D) conditions. In both conditions, Rpd3 is preferentially enriched at the promoters of these target genes, supporting a direct role in their transcriptional repression. (c) Western blot analysis of global H3K9ac levels in WT and *rpd3Δ* cells grown in +D and −D conditions. Loss of Rpd3 results in elevated H3K9ac levels, particularly upon glucose depletion, consistent with a role for Rpd3 as a histone deacetylase that removes activating acetyl marks during metabolic stress. Rpn10 was used as a loading control. (d) Box plots showing genome-wide H3K9ac enrichment (fold change over background) in WT and *rpd3Δ* strains under +D and −D conditions. Ordinary one-way ANOVA was used to calculate statistical significance. ****p<0.0001. The error bars depict the SEM. Strikingly, in both nutrient states, *rpd3Δ* cells exhibit lower H3K9ac enrichment compared to WT, despite a global increase in H3K9ac (from western blots). The reduced H3K9ac enrichment, perhaps, is indicative of more background acetylation in *rpd3Δ* cells. (e, f) Representative genome browser tracks displaying ChIP-seq signals for Rpd3 and H3K9ac at selected genes involved in fatty acid oxidation (FAO; e) and growth-related processes (f). These tracks illustrate the inverse relationship between Rpd3 occupancy and H3K9ac levels at specific loci. Arrows highlight regions where H3K9ac peaks are absent, and Rpd3 binding is high, emphasizing localized histone deacetylation associated with transcriptional repression. The H3K9ac peaks are retained in *rpd3Δ* cells.

To evaluate whether this binding influences promoter acetylation, we analyzed the distribution of H3K9ac in WT and *rpd3Δ* cells. Although immunoblotting revealed a global increase in H3K9ac levels upon Rpd3 loss [Figure 4c], ChIP–seq analysis showed a lower overall fold enrichment across the genome under both nutrient conditions, suggesting disrupted site-specific deacetylation and elevated background acetylation [Figure 4d]. This is consistent with a previous report showing that HDACs are required for limiting acetylation in gene bodies and intergenic regions for properly acetylated promoters^28^.

At Rpd3 target genes, however, the *rpd3Δ* strain displayed distinct promoter enrichment of H3K9ac. At the well-characterized Rpd3 target *INO1*, H3K9ac accumulation was evident at the promoter in *rpd3Δ* but absent in WT cells [Figure 4e]. A similar pattern was observed for two additional Rpd3-regulated genes active in +D—*CAT2* and *FOX2*—both involved in FAO, which aberrantly gained promoter H3K9ac in the *rpd3Δ* strain [Figure 4e]. Under –D conditions, Rpd3-target genes involved in growth and DNA replication, including *CLB5*, *CDC9*, and *DBF2*, also exhibited strong promoter H3K9ac enrichment in the *rpd3Δ* strain concomitant with transcriptional derepression [Figure 4f].

Collectively, Rpd3-bound genes could be divided into two categories. In one, Rpd3 deletion had a minimal effect on expression, corresponding to genes active under the current nutrient condition. In the other, Rpd3 occupancy was associated with repression of genes that are normally active under alternate metabolic states, indicating that Rpd3 could be pre-bound at their promoters to mediate nutrient-dependent repression during nutrient transitions.

### Rpd3–Gcn5 dynamics govern chromatin transitions during metabolic adaptation

Since histone acetylation at a given site depends on both HDAC as well as HAT activity, to examine how histone acetylation dynamics are coordinated during nutrient transitions, we next compared Rpd3 and Gcn5 occupancy under glucose-replete (+D) and glucose-depleted (–D) conditions. For growth-associated genes that failed to fully repress in the *rpd3Δ* strain during starvation, H3K9ac levels were high in +D but sharply decreased in –D in WT cells, consistent with their transcriptional status [Figure 5a]. Both Rpd3 and Gcn5 occupied these promoters, but their enrichment shifted reciprocally: Gcn5 binding declined while Rpd3 binding increased markedly upon glucose withdrawal [Figure 5b, c]. At the cell-cycle gene *CDC6*, for example, nutrient depletion led to reduced Gcn5 and elevated Rpd3 occupancy at the promoter in WT cells, coinciding with the loss of H3K9ac and transcriptional repression [Figure 5d]. The *rpd3Δ* strain, by contrast, retained promoter H3K9ac and displayed persistent transcriptional activity, as evident in the genome browser tracks [Figure 5d].

**Figure 5:**
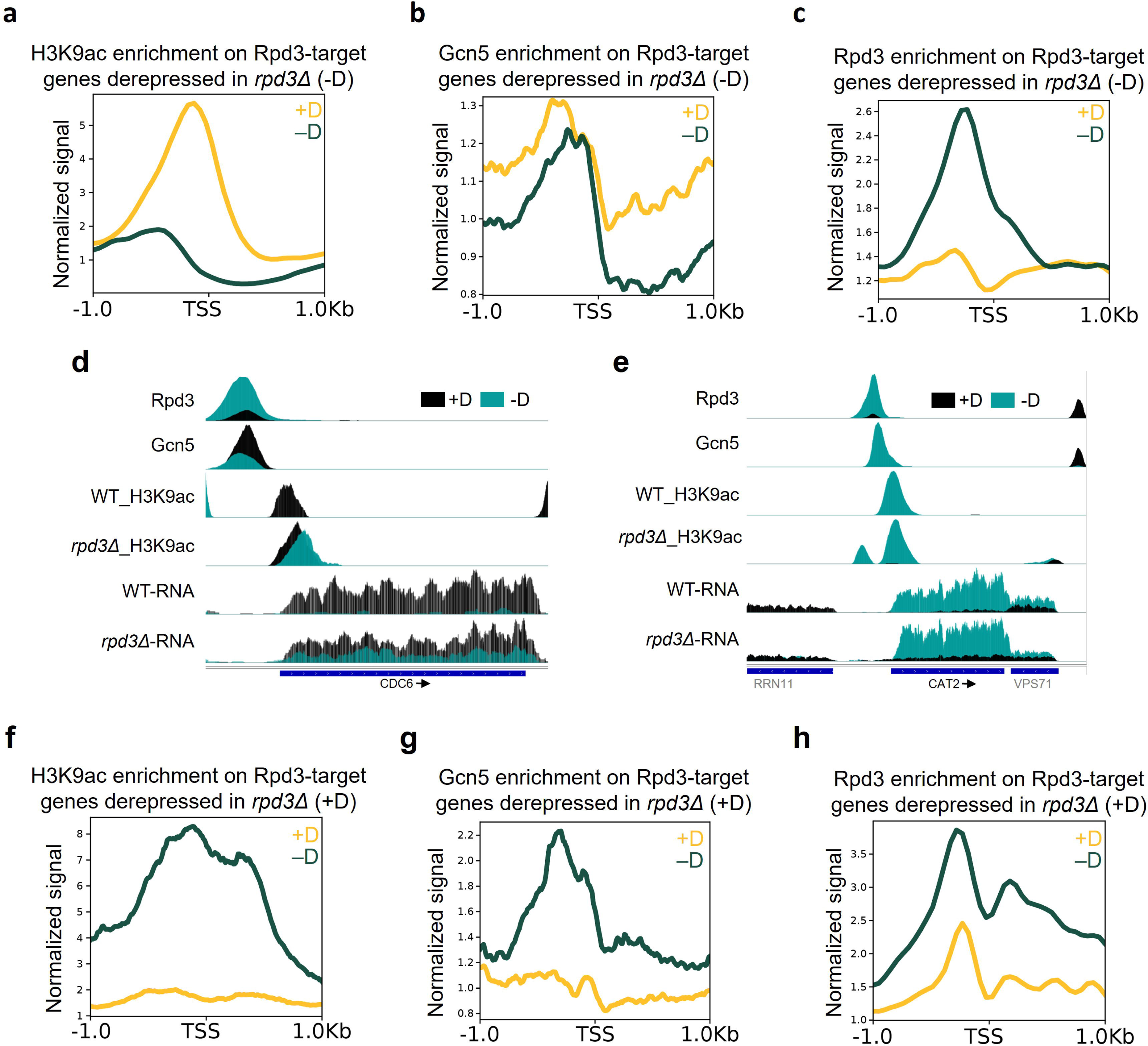
Dynamic interplay between Rpd3 and Gcn5 controls chromatin state during metabolic adaptation. (a-c, f-h) Metagene profiles derived from ChIP-seq datasets showing the genome-wide enrichment patterns of H3K9ac, the histone acetyltransferase Gcn5, and the histone deacetylase Rpd3 at genes that are derepressed upon *RPD3* deletion. Profiles are shown for cells grown under glucose-replete (+D) and glucose-depleted (−D) conditions. In both nutrient states, these genes exhibit coordinated occupancy of Gcn5 and Rpd3 along with altered H3K9ac levels, reflecting a dynamic balance between acetylation and deacetylation. Changes in the relative enrichment of these factors between +D and −D highlight condition-dependent chromatin remodeling associated with metabolic adaptation. (d, e) Representative genome browser tracks showing ChIP-seq signal for Rpd3, Gcn5, and H3K9ac, alongside RNA-seq read coverage (CPM-normalized), at selected loci. These examples illustrate the relationship between factor occupancy, histone acetylation, and transcriptional output. Regions with high Gcn5 binding and H3K9ac enrichment correspond to actively transcribed genes, whereas the correlation with Rpd3 occupancy is more nuanced. The deletion of *RPD3*, however, results in greater retention of H3K9ac, resulting in increased transcriptional output.

In contrast, genes involved in FAO that became derepressed in *rpd3Δ* cells under glucose conditions showed a different trend of Gcn5-Rpd3 binding dynamics. At these loci, including *CAT2*, both Gcn5 binding and H3K9ac signal were absent in +D, indicating complete disengagement of the acetyltransferase [Figure 5e]. This is consistent with their expression pattern in WT cells. This pattern was not unique to *CAT2* but extended to other Rpd3-target genes derepressed in *rpd3Δ* during +D [Figure 5f, g]. Unlike growth-related genes, FAO genes exhibited parallel enrichment patterns of Rpd3 and Gcn5, both reduced in +D. Under –D conditions, Rpd3 enrichment was more pronounced, suggesting a poised state consistent with its role in promoter deacetylation and transcriptional shutdown upon glucose repletion [Figure 5h].

Collectively, these analyses demonstrate that Rpd3 functions as a promoter-associated regulator that dynamically adapts to metabolic state. Growth-related genes display reciprocal Rpd3–Gcn5 modulation to fine-tune transcriptional output, whereas FAO genes follow a binary regulatory mode mediated by poised Rpd3 and Gcn5 disengagement. This coordinated interplay ensures precise, condition-specific transcriptional control during metabolic adaptation.

### Rpd3 exhibits nutrient-dependent redistribution across promoters and gene bodies

Unlike Gcn5, which primarily localizes to promoters, Rpd3 occupies both promoter regions and gene bodies [Figure 3]. Rpd3’s enrichment across gene bodies was markedly reduced upon nutrient depletion (–D) [Figure 3b], indicating that its chromatin association is dynamically remodeled during metabolic transitions.

Because Rpd3-mediated gene body deacetylation has been implicated in suppressing cryptic transcription^23,24^, which is associated with hallmarks of transcription such as increased chromatin accessibility and active histone marks^29^, we next examined whether Rpd3 enrichment at different gene elements correlated with transcription level. Indeed, Rpd3 was preferentially enriched across the coding regions of highly expressed genes compared with lowly expressed ones, consistent with its role in maintaining transcriptional fidelity at actively transcribed loci [Figure 6a, b].

**Figure 6:**
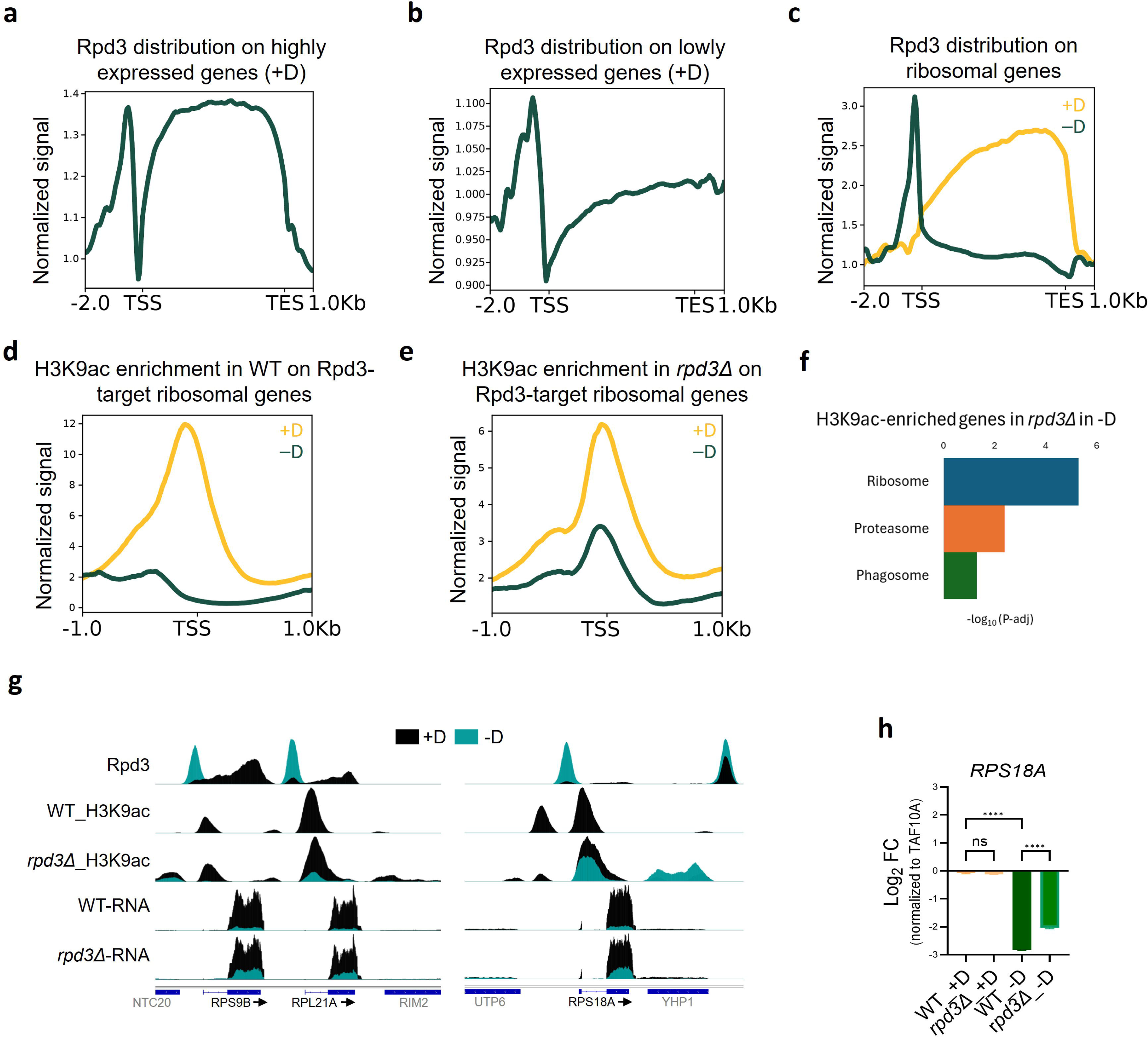
Rpd3 exhibits nutrient-dependent redistribution across gene elements to regulate transcription. (a-c) Metagene profiles illustrating the distribution of Rpd3 across gene elements (promoters, transcription start sites, and gene bodies) for the indicated gene categories under glucose-replete (+D) and glucose-depleted (−D) conditions. Rpd3 occupancy is markedly higher across gene bodies of highly expressed genes compared to lowly expressed genes in +D. Upon transition from +D to −D, Rpd3 undergoes a pronounced redistribution, shifting from gene bodies toward promoter regions, particularly at ribosomal protein genes, indicating a context-dependent change in chromatin engagement during metabolic adaptation. (d, e) Metagene plots showing H3K9ac enrichment centered at the transcription start sites (TSS) of Rpd3-bound ribosomal genes in +D and −D conditions. In −D, promoter-proximal accumulation of Rpd3 correlates with reduced H3K9ac levels, consistent with active histone deacetylation and transcriptional repression. In *rpd3Δ* cells, H3K9ac levels remain elevated under −D conditions, confirming that Rpd3 is required for deacetylation at these promoters during nutrient stress. (f) KEGG pathway enrichment analysis of genes that exhibit increased H3K9ac levels in *rpd3Δ* cells under glucose-depleted (−D) conditions. Enriched pathways prominently include ribosome biogenesis and translation-related processes, indicating that loss of Rpd3 impairs repression of growth-associated gene programs during metabolic stress. (g) Representative genome browser tracks displaying ChIP-seq signal for Rpd3 and H3K9ac, along with RNA-seq read coverage (CPM-normalized), at selected loci in WT and *rpd3Δ* cells. These examples demonstrate the redistribution of Rpd3 from gene bodies to promoter regions upon glucose depletion, accompanied by a decrease in H3K9ac levels and reduced transcription in WT cells. In contrast, *rpd3Δ* cells show sustained H3K9ac enrichment and attenuated transcriptional repression, consistent with the metagene analyses. (h) qPCR analysis of the representative ribosomal protein gene *RPS18A* in WT and *rpd3Δ* strains under +D and −D conditions. Expression levels were measured relative to an internal control, *TAF10A*, and reveal that Rpd3 deletion results in impaired downregulation of *RPS18A* upon glucose depletion, supporting a role for Rpd3 in repressing ribosomal gene expression during nutrient limitation. Ordinary one-way ANOVA was used to calculate statistical significance. n=3, ****p<0.0001. The error bars depict the SEM.

Rpd3 binding in both nutrient states was strongly enriched at ribosomal genes [Figure 1d]. As ribosomal genes are among the most highly expressed under glucose-replete conditions (+D) but are strongly repressed upon starvation (–D) [Supplementary Figure S1a, b, d], we next analyzed Rpd3 occupancy at these loci. Under +D conditions, Rpd3 was distributed across ribosomal gene bodies [Figure 6c]. Upon glucose depletion, however, its binding shifted sharply toward promoter regions, coinciding with transcriptional repression of these genes [Figure 6c]. ChIP–seq analysis of H3K9ac revealed an inverse pattern: acetylation near the TSSs decreased where Rpd3 binding increased, showing a clear anticorrelation between promoter recruitment of Rpd3 and histone acetylation [Figure 6d].

To determine whether Rpd3 directly regulates promoter acetylation and expression of ribosomal genes, we analyzed H3K9ac profiles and transcript levels in the *rpd3Δ* strain. In contrast to WT cells, where H3K9ac is depleted from ribosomal promoters during starvation, the *rpd3Δ* strain partially retained promoter H3K9ac enrichment under the same conditions [Figure 6d, e]. Pathway analysis of H3K9ac-enriched genes confirmed that H3K9ac remained concentrated at ribosomal loci [Figure 6f], rather than redistributing to respiration-associated genes as in WT cells [Figure 1a]. These global patterns were evident in genome browser tracks of representative ribosomal genes, including *RPL21A* and *RPS18A* [Figure 6g]. Consistent with these observations, *RPS18A* transcript levels remained elevated in *rpd3Δ*, as validated by qPCR [Figure 6h]. Notably, Rpd3 has been reported to be involved in the regulation of rRNA genes as well^30^.

Together, these results demonstrate that Rpd3’s chromatin occupancy can be nutrient dependent even on a given gene. Under glucose-rich conditions, Rpd3 associates with gene bodies of highly expressed genes, perhaps, to preserve transcriptional fidelity, whereas during starvation, its redistribution to promoters mediates targeted deacetylation and repression of growth-associated transcriptional programs.

### Rpd3’s genome-wide distribution is shaped by its association with complex-specific subunits

Since Rpd3 binds both promoter and gene body regions and undergoes redistribution upon nutrient shifts, we next examined how this transition is orchestrated. Rpd3 functions within at least two major histone deacetylase complexes, Rpd3L and Rpd3S, which share the core subunits Rpd3, Sin3, and Ume1 but differ in their auxiliary components that dictate chromatin targeting and function. The large complex (Rpd3L) contains Pho23, a subunit linked to promoter recruitment through recognition of H3K4me3, while the small complex (Rpd3S) includes Eaf3, which recognizes H3K36me3-decorated gene bodies to suppress cryptic transcription^31,32^.

To determine how these complexes contribute to Rpd3 recruitment, we performed ChIP–seq for Pho23 (Rpd3L) and Eaf3 (Rpd3S) in cells grown in glucose (+D) and after 30 minutes of glucose depletion (–D). Under glucose-replete conditions, most Pho23- and Eaf3-bound genes overlapped with Rpd3-bound loci, consistent with their physical association in Rpd3-containing complexes [Figure 7a]. However, distinct subsets of genes were co-bound by Rpd3 and either Eaf3 or Pho23, indicating complex-specific targeting [Figure 7a]. Strikingly, the majority of Rpd3-bound genes were associated with both subunits, suggesting that Rpd3L and Rpd3S may not be entirely mutually exclusive *in vivo* and could coexist or dynamically interchange at certain loci. This was reflected in their binding distributions as well: Pho23 displayed stronger promoter enrichment, while Eaf3 was slightly more prominent across gene bodies, especially at the 3’ end; nonetheless, both subunits showed occupancy at both regions [Supplementary figure S4a].

**Figure 7:**
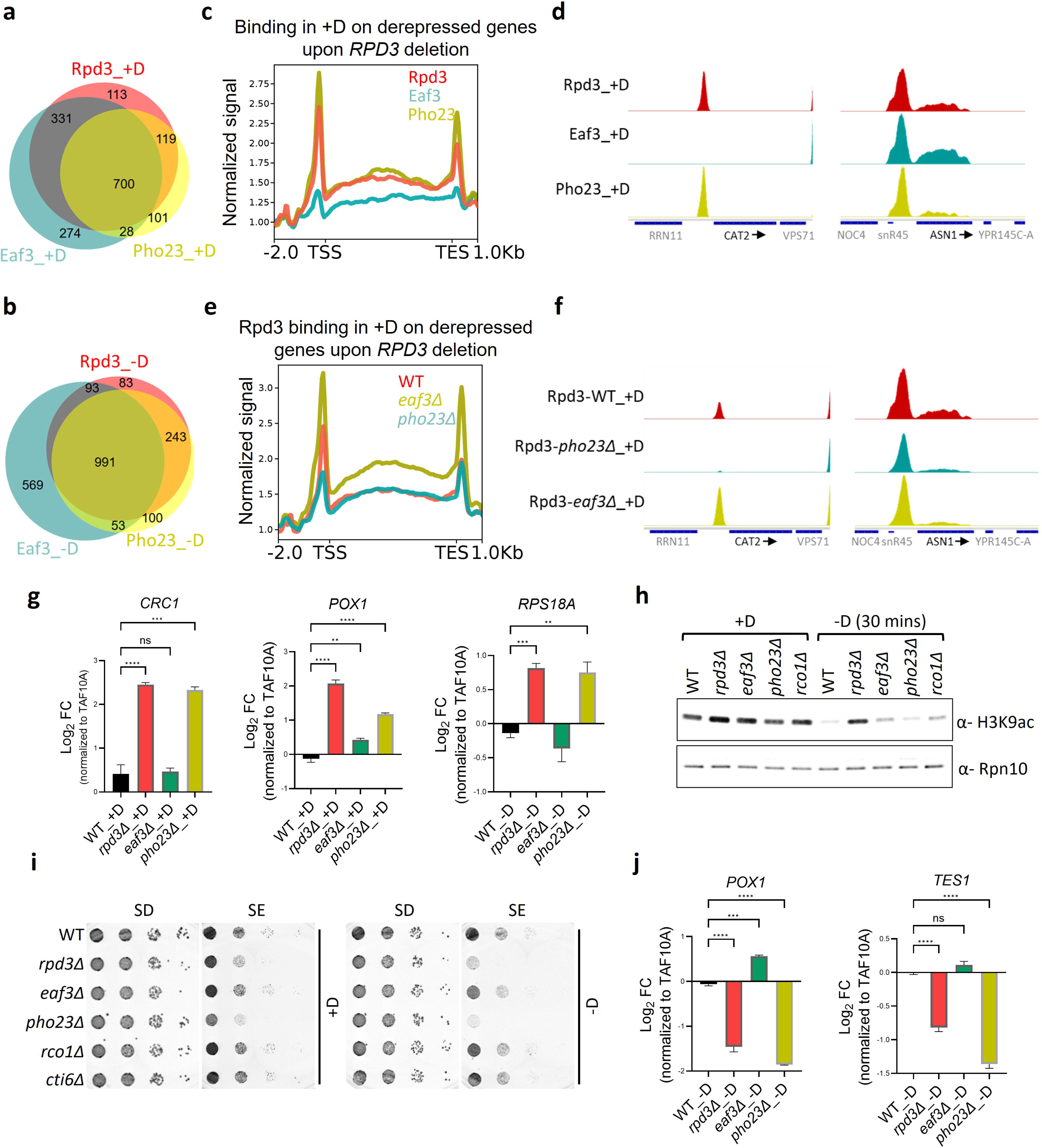
The large complex mediates the promoter recruitment of Rpd3 to nutrient-regulated genes. (a, b) Venn diagrams illustrating the overlap of genomic targets bound by Rpd3, Eaf3 (a component of the Rpd3S complex), and Pho23 (a subunit of the Rpd3L complex) in glucose-replete (+D) and glucose-depleted (−D) conditions. Interestingly, a substantial overlap exists between Pho23- and Eaf3-bound genes, which suggests that the Rpd3L and S complexes may have overlaps. (c) Metagene profiles showing the genome-wide distribution of Rpd3, Eaf3, and Pho23 specifically at genes that are derepressed upon *RPD3* deletion in +D. Pho23 exhibits strong enrichment at promoter regions, consistent with its proposed role in directing Rpd3L to promoter-proximal chromatin. In contrast, Eaf3 shows relatively low enrichment at these sites, indicating that the Rpd3S complex plays a limited role in Rpd3 recruitment to the majority of derepressed genes. (d) Representative genome browser tracks displaying ChIP-seq peaks for Rpd3, Eaf3, and Pho23 at selected loci. Eaf3 binding is notably reduced or absent at these genes, further supporting the idea that Rpd3S is not the primary recruiter of Rpd3 at these sites. In contrast, Pho23 shows robust co-occupancy with Rpd3, particularly at promoter regions, underscoring the role of the Rpd3L complex in chromatin targeting. (e) Metagene plot comparing Rpd3 occupancy at genes derepressed upon *RPD3* deletion in WT, *eaf3Δ*, and *pho23Δ* strains grown under +D conditions. Deletion of *PHO23* results in a marked reduction of Rpd3 binding at promoters of these genes, whereas loss of *EAF3* has minimal impact. This demonstrates that Pho23, and by extension the Rpd3L complex, is essential for recruiting Rpd3 to its regulatory targets under nutrient-replete conditions. (f) Representative genome browser tracks showing Rpd3 ChIP-seq signal in WT, *eaf3Δ*, and *pho23Δ* cells at selected loci under +D conditions. Deletion of *PHO23* causes a pronounced loss of Rpd3 binding at metabolic genes such as the fatty acid oxidation gene *CAT2*, whereas Rpd3 occupancy at a control gene (e.g., *ASN1*) remains unchanged. This highlights the gene-specific requirement for Pho23 in mediating Rpd3 recruitment, particularly at nutrient-responsive loci. (g) qPCR analysis of representative fatty acid oxidation (FAO) genes and growth-related genes in WT, *rpd3Δ*, *eaf3Δ*, and *pho23Δ* strains grown under glucose-replete (+D) or glucose-depleted (−D) conditions. Gene expression levels were normalized to an internal control, *TAF10A*. Ordinary one-way ANOVA was used to calculate statistical significance. n=3, ns>0.9999, **p=0.0059, ***p=0.0001, ****p<0.0001. The error bars depict the SEM. In both nutrient states, the *pho23Δ* strain exhibits elevated transcript levels comparable to those observed in *rpd3Δ* cells, indicating that the Rpd3L complex is required for repression of these genes. In contrast, deletion of *EAF3*, a subunit of the Rpd3S complex, does not significantly alter expression of these targets, demonstrating functional specificity of the Rpd3L complex in regulating nutrient-responsive genes. (h) Western blot analysis of global H3K9ac and total H3 acetylation (H3ac) levels in WT and mutant strains. Cells were grown in glucose-replete (+D) media to mid-exponential phase (OD600 ≈ 1), followed by a shift to glucose-depleted (−D) conditions for 30 minutes. Whole-cell lysates were prepared and analyzed by western blot, with Rpn10 serving as a loading control. Deletion of *EAF3* or *RCO1*, both components of the Rpd3S complex, results in greater retention of H3K9ac upon glucose depletion compared to WT, indicating that the Rpd3S complex is required for the global histone deacetylation response during metabolic stress. In contrast, *pho23Δ* shows a phenotype more similar to WT, suggesting that Rpd3L plays a more gene-specific role, whereas Rpd3S mediates genome-wide deacetylation. (i) Growth spotting assay assessing the ability of WT and mutant strains to adapt to nutrient limitation. Cells were grown in +D to mid-exponential phase (OD600 ≈ 1), shifted to −D for 30 minutes, and then serially diluted and spotted onto synthetic dextrose (SD; glucose-containing) and synthetic ethanol (SE; non-fermentable carbon source) agar plates. Images have been color-inverted for improved visualization. The *pho23Δ* strain exhibits a growth defect on SE plates comparable to that of *rpd3Δ*, indicating that the Rpd3L complex is essential for metabolic adaptation and growth on non-fermentable carbon sources. In contrast, *eaf3Δ* shows growth similar to WT, further supporting the distinct and non-redundant roles of Rpd3L and Rpd3S complexes. (j) qPCR analysis of representative FAO genes in WT, *rpd3Δ*, *eaf3Δ*, and *pho23Δ* strains under glucose-depleted (−D) conditions. Ordinary one-way ANOVA was used to calculate statistical significance. n=3, ns>0.9999, ***p=0.0001, ****p<0.0001. The error bars depict the SEM. Consistent with results from +D (panel a), *pho23Δ* cells display elevated expression of FAO genes similar to *rpd3Δ*, while *eaf3Δ* shows minimal derepression. This confirms that the Rpd3L complex, via Pho23, is the primary mediator of Rpd3-dependent transcriptional repression at metabolic genes under both nutrient conditions, whereas Rpd3S functions predominantly in global chromatin maintenance rather than gene-specific regulation.

Upon glucose depletion, a marked redistribution was observed. The overlap between Pho23- and Rpd3-bound loci expanded substantially, while the overlap between Eaf3 and Rpd3 decreased [Figure 7b]. Binding profiles showed enhanced Pho23 enrichment at promoters and Eaf3 occupancy across gene bodies [Supplementary figure S4b]. These shifts indicate that under starvation, the promoter-directed Rpd3L complex plays a more pronounced role in Rpd3 recruitment as compared to glucose, consistent with its increased promoter occupancy during nutrient stress.

To directly test the contribution of each subunit to Rpd3 recruitment, we performed Rpd3 ChIP–seq in *pho23Δ* and *eaf3Δ* strains. Unexpectedly, deletion of Eaf3 resulted in increased promoter enrichment of Rpd3, with minimal changes across gene bodies [Supplementary Figure S4c, d], suggesting that the absence of Eaf3 may result in a greater recruitment of Rpd3 to the promoters, possibly by the large complex. In contrast, Pho23 deletion caused only a modest reduction in Rpd3 occupancy, indicating potential redundancy or alternative pathways for promoter recruitment. Also, the deletion of a subunit may impact the complex’s stoichiometry and stability, resulting in its aberrant recruitment.

These findings reveal that Rpd3’s genome-wide distribution is shaped by the dynamic balance between Rpd3L and Rpd3S complex associations. The large and small complexes appear to cooperate rather than function in strict isolation, enabling Rpd3 to reallocate between promoter and gene body chromatin in response to changing metabolic conditions. Nutrient shifts remodel this equilibrium—favoring Rpd3S-mediated deacetylation of coding regions under growth conditions and Rpd3L-mediated promoter repression during starvation.

### The Rpd3L subunit Pho23 mediates promoter recruitment of Rpd3 to nutrient-regulated genes

The genome-wide binding profiles of Rpd3 complex-specific subunits and their role in Rpd3’s recruitment uncovered correlations but also revealed nuances suggestive of complex interplay between regulatory mechanisms. Rpd3’s role in genome-wide regulation of transcription may involve the site-specific utilization of certain complex components^21^. To test how the Rpd3L and Rpd3S complexes contribute to transcriptional regulation of Rpd3 target genes that are specifically derepressed in the *rpd3Δ* strain, we analyzed Pho23 and Eaf3 binding at these genes.

At FAO genes that are inappropriately expressed in *rpd3Δ* during glucose growth (+D), Pho23—but not Eaf3—was strongly enriched at promoters [Figure 7c]. A similar enrichment pattern was observed at ribosomal genes that fail to fully repress in *rpd3Δ* under starvation (–D) [Supplementary figure S4e]. This promoter-proximal localization of Pho23 was clearly visible in genome browser tracks for representative loci such as *INO1* [Supplementary figure S4f], *CAT2* (FAO gene, +D) [Figure 7d], and *RPS9B*, *RPL21A*, and *CDC6* (growth-associated genes, –D) [Supplementary figure S4g]. In contrast, genes that remain unaffected by nutrient transitions but are Rpd3-bound, such as *ASN1*, show similar enrichment of both Pho23 and Eaf3 [Figure 7d].

To determine whether Pho23 directly mediates Rpd3 recruitment at these genes, we checked Rpd3’s binding in *pho23Δ* and *eaf3Δ* strains. Loss of Pho23 caused a marked reduction in Rpd3 occupancy at promoters of the *INO1* gene [Supplementary Figure S4h] as well as these nutrient-regulated genes in both +D and -D [Figure 7e, supplementary figure S4i]. Consistent with our earlier observation of the genome-wide binding pattern, deletion of Eaf3 resulted in increased promoter binding of Rpd3 [Figure 7e, supplementary figure S4i]. These outcomes were evident at representative loci, where Rpd3 peaks were substantially diminished in *pho23Δ* cells both at *CAT2* in +D and *RPS18A* in –D [Figure 7f, supplementary figure S4j]. In contrast, Rpd3 occupancy at genes not responsive to nutrient shifts, such as *ASN1*, remained largely unchanged in the subunit-specific deletion strains [Figure 7f].

Together, these findings demonstrate that Pho23 directs Rpd3 recruitment to promoters of genes that require repression during metabolic transitions. This promoter-specific recruitment enables Rpd3 to impose histone deacetylation and transcriptional silencing in response to carbon source changes.

### Distinct roles of Rpd3L and Rpd3S complexes in global and promoter-specific deacetylation and metabolic adaptation

Our results indicated that Pho23 recruits Rpd3L to promoters of nutrient-responsive genes, while Eaf3 mediates Rpd3S association with gene bodies. To determine how each complex contributes functionally to chromatin and transcriptional changes observed upon Rpd3 deletion, we analyzed the effects of Pho23 and Eaf3 loss on gene expression and histone acetylation.

Consistent with its role in promoter-specific recruitment, Pho23 deletion resulted in derepression of Rpd3 target genes, similar to the phenotype of *rpd3Δ*. Genes such as *CRC1*, *POX1*, *CAT2*, *FOX2*, and *TES1* (derepressed in +D), as well as *RPS18A* and *RPL21A* (upregulated in –D), exhibited increased expression in *pho23Δ* but not in *eaf3Δ* [Figure 7g, supplementary figure S5a]. This indicates that Rpd3L, but not Rpd3S, is required for transcriptional repression of these nutrient-regulated genes.

We next examined genome-wide H3K9ac levels in *pho23Δ* and *eaf3Δ* strains under both nutrient conditions. Pho23 deletion did not noticeably alter global acetylation dynamics relative to WT, whereas Eaf3 deletion caused a small but consistent retention of H3K9ac during both +D and –D conditions [Figure 7h]. Deletion of another Rpd3S-specific subunit, Rco1, produced an identical effect on global H3K9ac, confirming that Rpd3S at least partly mediates the broad deacetylation observed during nutrient shifts [Figure 7h]. This is consistent with previous observations that, in addition to targeted recruitment, Rpd3 also deacetylates large regions of chromatin, including coding regions^33^.

To determine how these chromatin changes translate into physiological outcomes, we compared the ability of WT, *rpd3Δ*, *pho23Δ*, and *eaf3Δ* cells to grow in respiratory media. Cells were first grown in a synthetic media containing glucose (SD) and then shifted to ethanol (SE) as a non-fermentable carbon source. Consistent with their similar gene expression profiles, both the *rpd3Δ* and *pho23Δ* strains exhibited impaired growth under respiratory conditions, irrespective of whether they were pre-grown in +D or -D, whereas *eaf3Δ* cells grew comparably to WT [Figure 7i]. This defect correlated with reduced induction of respiration-associated genes, including *POX1* and *TES1*, in *rpd3Δ* and *pho23Δ* mutants [Figure 7j].

These findings establish a division of labor between the two Rpd3 complexes: Rpd3S mediates global histone deacetylation, whereas Rpd3L executes promoter-specific repression through Pho23 recruitment. Together, these mechanisms ensure proper transcriptional and metabolic adaptation during transitions in carbon sources.

## DISCUSSION

Our findings demonstrate that promoter-poised deacetylation by an HDAC complex is central to maintaining transcriptional fidelity during nutrient transitions in *Saccharomyces cerevisiae*. Rpd3 dynamically redistributes across chromatin in response to metabolic cues, implementing both local and global deacetylation via its two complexes, Rpd3L and Rpd3S. Rpd3L—anchored at promoters via its subunit Pho23—enables rapid histone deacetylation and transcriptional shutdown when nutrient availability changes. In parallel, Rpd3S—directed by Eaf3 and Rco1—ensures gene-body deacetylation, perhaps to prevent aberrant or cryptic transcription. Together, these mechanisms coordinate a precise and reversible chromatin response enabling yeast to switch between fermentative and respiratory growth programs.

### Resolution of the HDAC paradox

Multiple genome-wide studies have noted that HDACs, including Rpd3, are paradoxically enriched at active promoters despite their canonical role as repressors^20–22^. A study in flies suggested that HDACs are required for promoter-proximal Pol II pausing^34^. Since yeast cells lack Pol II pausing, this model doesn’t explain the counterintuitive observation of HDAC presence at yeast promoters. Our data resolve this paradox by showing that Rpd3’s promoter occupancy at active genes reflects a poised regulatory state rather than constitutive repression. By remaining bound at promoters of actively transcribed genes via Rpd3L, the complex allows cells to rapidly repress these genes in response to nutrient perturbation. This model aligns with the notion that chromatin regulators act as “epigenetic toggles,” occupying key genomic positions to facilitate rapid transcriptional switching in response to environmental changes, instead of affecting the steady-state level of transcripts that are regulated primarily by transcription factors^35–37^. These toggles function within a broader chromatin framework rather than in isolation. As recently shown, the combined influence of DNA sequence, genomic context, and interacting histone modifications determines how transcriptional responses are modulated^38^. Consequently, chromatin marks serve not as solitary determinants of gene expression but as components of a multilayered and context-dependent regulatory system that orchestrates dynamic transcriptional control.

### Rpd3L as a chromatin memory module

The promoter-associated Rpd3L complex has been previously implicated in transcriptional repression memory (TREM), where transient repression is stably inherited through histone deacetylation linked to the H3K4me3 mark^25^. Our results extend this model by demonstrating that Rpd3L not only maintains repression memory but also acts dynamically to modulate transcription during metabolic shifts. The Pho23 subunit of the Rpd3L complex is especially important for this since the deletion of a different subunit, Cti6, did not result in a phenotype in SE [Figure 7i]. This is consistent with previous findings that resulted in the conclusion that even though Pho23 and Cti6 bind to the same histone mark, they may have distinct functions in transcription^25^. The enrichment of Pho23-bound Rpd3L at promoters of nutrient-responsive genes suggests that Rpd3L senses promoter context likely through H3K4me3 and transcription factor interactions, allowing targeted repression when transcriptional programs must be silenced. Thus, Rpd3L appears to fulfill a dual role—both establishing chromatin memory and enabling adaptive regulation of transcription.

### Rpd3S ensures transcriptional fidelity at active genes

In parallel with the promoter-directed functions of Rpd3L, the Rpd3S complex contributes to global chromatin homeostasis by deacetylating the gene bodies of highly expressed genes. This activity at least partly relies on the Eaf3 (and Rco1) subunit, which recognizes the H3K36me3 mark deposited co-transcriptionally on active genes, consistent with previous findings that Rpd3S suppresses cryptic transcription initiation within coding regions^23,39^. In line with this, deletion of Rpd3S-specific subunits led to a modest retention of global H3K9ac signal [Figure 7h] but did not result in promoter derepression [Figure 7g, supplementary figure 5a], underscoring the complex’s specialized role in maintaining transcriptional fidelity rather than directly regulating canonical promoter output.

Notably, during starvation, the promoter-directed Rpd3L complex appears to contribute more prominently to Rpd3 recruitment than under glucose-replete conditions, as evidenced by the increased overlap between Rpd3 and Pho23 occupancy in –D [Figure 7a, b]. This shift suggests enhanced engagement of the Rpd3L complex at promoters during nutrient depletion. Importantly, transcriptional profiling revealed no significant changes in the expression of Rpd3L or Rpd3S subunits upon transition from +D to –D, indicating that altered recruitment is unlikely to result from changes in subunit abundance [Table S1]. Instead, these observations raise the possibility that nutrient-dependent remodeling of complex composition or post-translational modifications (PTMs) within Rpd3-containing complexes may modulate their chromatin association. Such mechanisms could dynamically tune Rpd3 targeting without requiring changes in gene expression of its constituent subunits that are often shared with other multiprotein complexes.

Nevertheless, the division of labor between Rpd3L and Rpd3S provides a mechanistic explanation for how a single histone deacetylase can coordinate both localized and global chromatin responses to metabolic cues. While Rpd3L mediates targeted promoter repression during nutrient transitions, Rpd3S safeguards the transcriptional fidelity of actively transcribed regions.

### Histone deacetylation may replenish the acetate pool

Our observation that global histone deacetylation accompanies nutrient shift—and that this activity is largely diminished in *rpd3Δ* cells—raises the possibility that active deacetylation serves a metabolic recycling function. By removing acetyl groups from histones, Rpd3 may liberate acetate, which can be converted via the nuclear acetyl-CoA synthetase Acs2 into acetyl-CoA, thereby feeding HAT-mediated reacetylation of promoters required for the starvation response. This model is consistent with recent reports linking chromatin-bound acetyl pools to metabolic flux^40^. Such a mechanism would be particularly relevant in –D conditions, where acetate availability is limited and “trapped” acetyl groups may serve as a metabolic reserve. However, despite this proposed role, deletion of Rpd3S components did not cause overt growth defects in starvation media (SE). It is possible that the individual deletion of Eaf3 or Rco1 [Figure 7h], or their combined loss [Supplementary Figure S5b], did not substantially diminish global Rpd3S deacetylase activity, and this may contribute to the lack of phenotype. Another possibility is that additional deacetylases, such as Hda1, may need to be deleted in conjunction with Rpd3 to observe the full extent of phenotypes. A previous study demonstrated that complete derepression of the Rpd3 target *PHO5* occurred only in the *hda1Δ rpd3Δ* double mutant^33^, suggesting that although Rpd3 plays a primary role as the histone deacetylase in nutrient-dependent regulation, it may act cooperatively with other HDACs. Consistent with this notion, our western blot analysis showed that Hda1 deletion also led to detectable retention of histone acetylation under –D conditions [Figure 1c], supporting the idea of partial redundancy among HDACs in maintaining chromatin deacetylation during metabolic transitions.

### Coordination between Gcn5 and Rpd3 during nutrient transitions

The extensive Rpd3-mediated deacetylation of histones raises an important mechanistic question: what triggers this response? Do fluctuations in glucose availability directly modulate Rpd3 catalytic activity, or is Rpd3 actively recruited to specific target genes that require deacetylation during nutrient transitions? Although these possibilities are not mutually exclusive, our data indicates that the net deacetylation observed under glucose-depleted (–D) conditions arises from a finely tuned coordination between opposing acetylation and deacetylation machineries. Previously, we have shown that Gcn5 undergoes transcription factor-driven chromatin recruitment and redistribution upon the +D to -D switch^19^. Since widespread co-occupancy of Rpd3 and Gcn5 exists at active promoters, once Gcn5 disengages a locus, triggered by nutrient-dependent redistribution, the action of Rpd3 results in the net loss of acetylation at those sites, resulting in gene repression.

Interestingly, the dynamics of Gcn5 and Rpd3 binding differ depending on the functional class of the target genes. At growth-related genes, Gcn5 is bound under both glucose-replete (+D) and glucose-depleted (–D) conditions, with its enrichment correlating positively with the transcriptional demand of these genes [Figure 5b]. This pattern is consistent with the requirement of growth programs across nutrient states, albeit at varying levels. Since Gcn5 occupancy remains relatively stable, the increase in Rpd3 binding observed upon starvation results in a net loss of H3K9ac, leading to transcriptional repression [Figure 5c].

In contrast, at FAO genes, Gcn5 exhibits a binary binding mode—present under starvation (–D) but absent in glucose-rich conditions (+D) [Figure 5g]. This binary switch reflects the conditional activation of these genes, which are unnecessary during glycolytic growth. Correspondingly, Rpd3 binding is markedly reduced in +D, likely due to the diminished requirement for HDAC activity in the absence of a recruited HAT [Figure 5h].

The outcome of this reciprocal Rpd3–Gcn5 modulation is pathway-dependent: for growth genes, their dynamic interplay fine-tunes transcriptional amplitude, whereas for FAO genes, it enforces rapid ON/OFF switching in response to nutrient cues. The molecular determinants governing these differential enrichment patterns remain unclear. This suggests that transcription factor recruitment, post-translational modifications, or crosstalk with other histone marks may drive these context-specific interactions, which will be interesting to study in the future.

While Gcn5 promotes H3K9ac deposition and transcriptional activation, Rpd3 ensures timely deacetylation and repression upon metabolic shift. This antagonistic interplay exemplifies a chromatin-based feedback mechanism through which histone marks are dynamically reset, allowing transcriptional programs to remain responsive to the cell’s metabolic state rather than locked in static ON or OFF configurations.

### Metabolic control of chromatin dynamics

The coupling of Rpd3 activity to metabolic state underscores the central role of chromatin in translating environmental inputs into transcriptional outcomes. By mediating reversible deacetylation, Rpd3 provides an epigenetic checkpoint that prevents inappropriate persistence of growth programs while enabling activation of catabolic pathways during starvation. This regulatory framework echoes broader principles observed in more complex eukaryotes, where HDAC complexes integrate metabolic and signaling pathways to maintain transcriptional homeostasis^41,42^.

In conclusion, our study defines Rpd3L as a promoter-poised HDAC that governs transcriptional fidelity during metabolic transitions by dynamically redistributing across chromatin. Through coordinated action with Gcn5 and the Rpd3S complex, Rpd3L ensures rapid, reversible, and accurate transcriptional reprogramming in response to nutrient changes. These findings bridge metabolism and chromatin regulation, offering a unified model for how cells integrate environmental signals into precise transcriptional outputs.

#### Limitation of the study

While our study highlights the role of HDACs in reshaping the transcriptional landscape during cellular adaptation to nutrient shifts, several questions remain unresolved. In particular, the direct impact of nutrient availability on post-translational modifications (PTMs), enzymatic activity, and the composition of HDAC-containing complexes was not examined and warrants further investigation. Additionally, although the Rpd3S complex is well established in suppressing cryptic transcription, we did not directly assess the effect of Rpd3 complex mutants on cryptic transcript abundance. Given that Rpd3S appears to contribute significantly to the widespread deacetylation observed following nutrient switching, this represents an important area for future study.

## RESOURCE AVAILABILITY

### Lead contact

Requests for further information and resources should be directed to the lead contact, Benjamin P. Tu.

### Materials availability

All unique/stable reagents generated in this study are available from the lead contact without restriction.

### Data and code availability

- ChIP- and RNA-Seq datasets are available in the Gene Expression Omnibus (GEO) database under the accession numbers GSE178160 (RNA-seq), GSE178159 (H3K9ac, H3, Gcn5-HA ChIP-Seq) and GSE312645 (Rpd3-HA, Pho23-HA, Eaf3-HA ChIP-Seq) and are publicly available as of the date of publication.
- This paper does not report original code.
- Any additional information required to reanalyze the data reported in this paper is available from the lead contact upon request.

## ACKNOWLEDGMENTS

The authors are grateful to the Tu lab members for their useful discussion. Additionally, we thank Dr. Divya Reddy of UTSW, Dallas, Texas, USA, for her constructive feedback on the manuscript. This work was supported by NIH grants R35GM136370, R01NS115546, and HHMI.

## AUTHOR CONTRIBUTIONS

Conceptualization, S.B.; methodology, S.B. and B.M.S; Investigation, S.B. and B.M.S; writing – original draft, S.B.; writing – review & editing, S.B. and B.P.T.; funding acquisition, B.P.T; supervision, B.P.T.

## SUPPLEMENTAL INFORMATION

Document S1. Figures S1–S5, Table S2

Table S1. Differential gene expression analysis derived from RNA-Seq data

## STAR★METHODS

### KEY RESOURCES TABLE

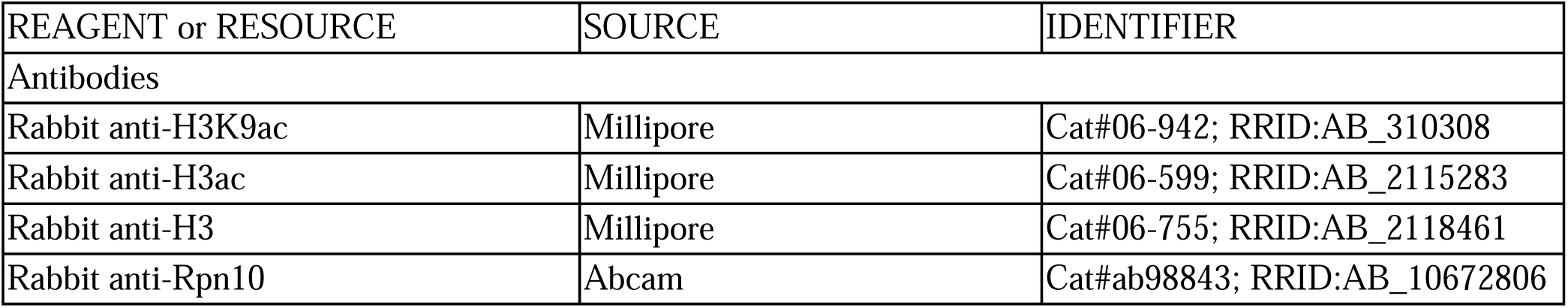
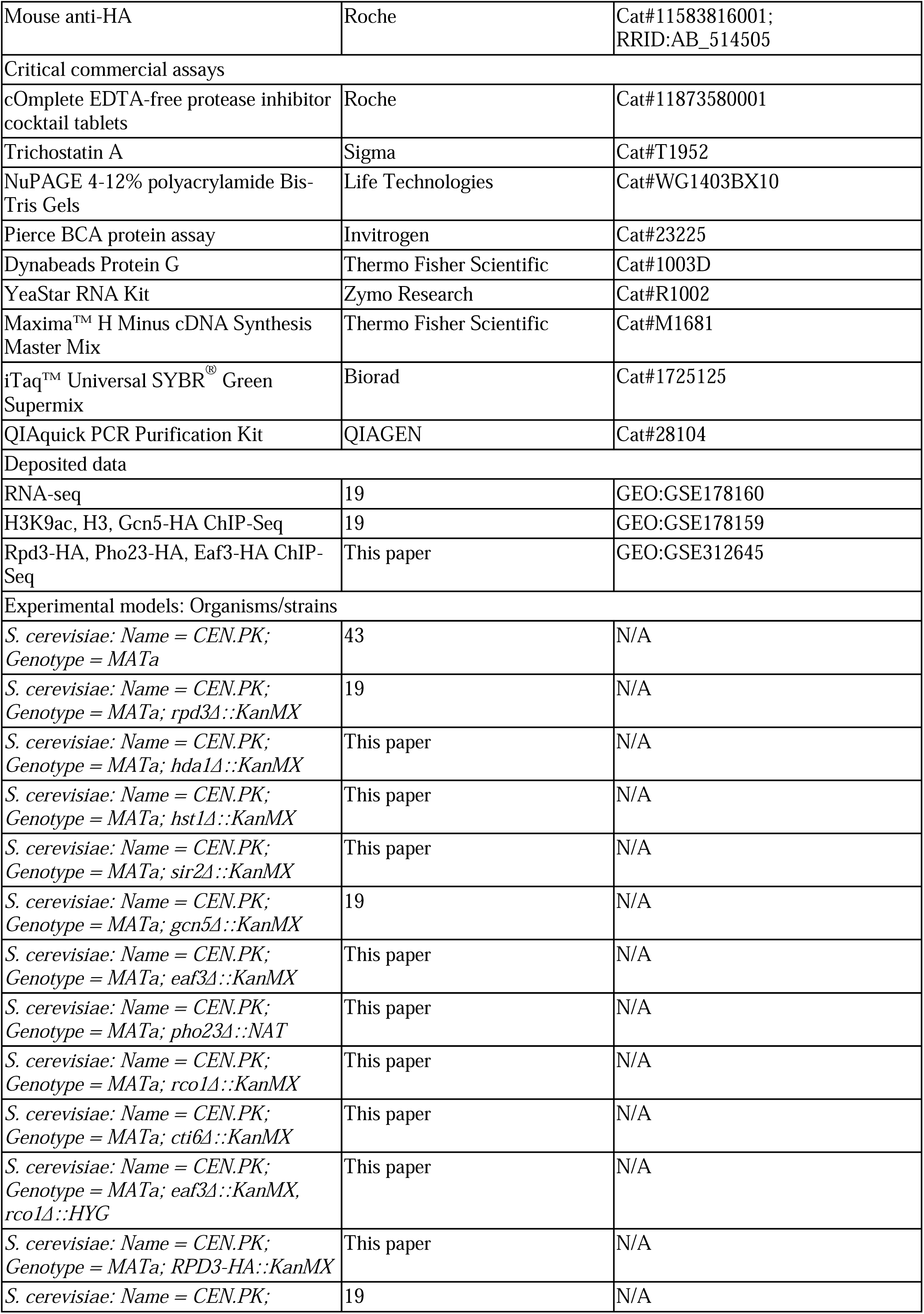
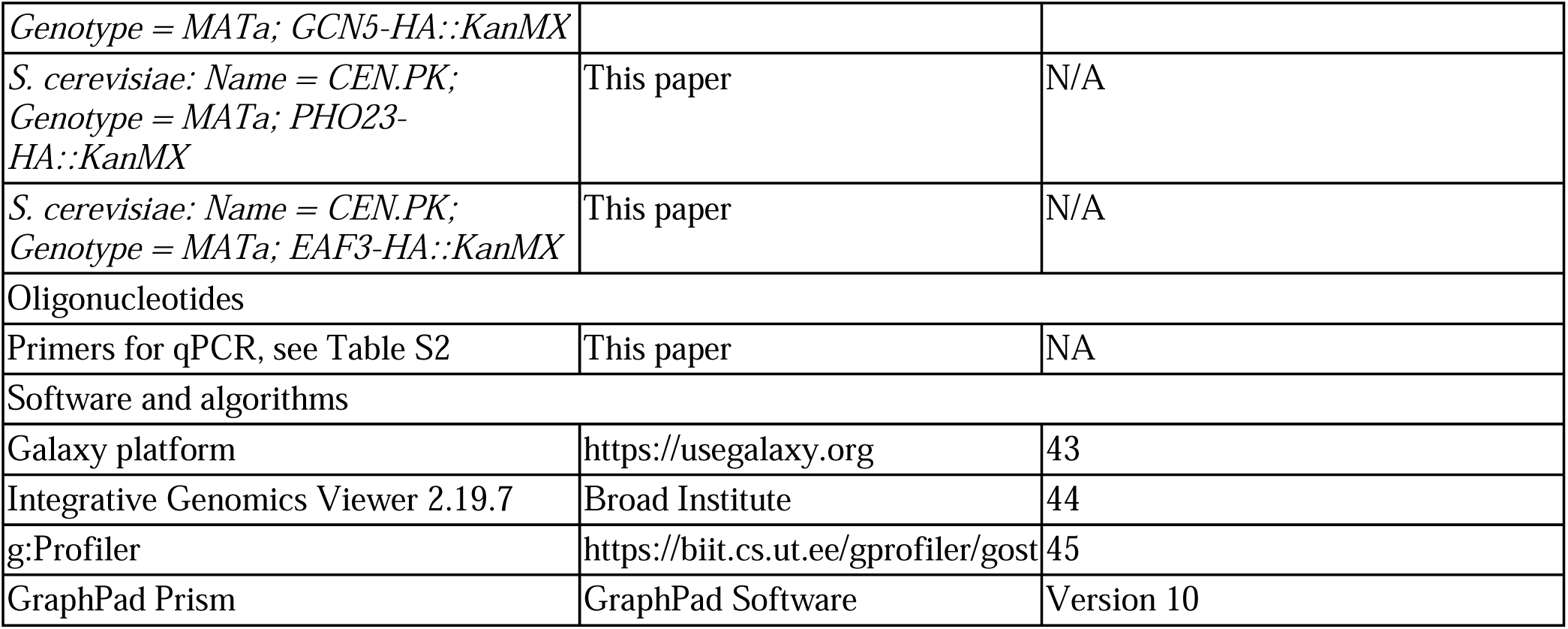

### EXPERIMENTAL MODEL AND STUDY PARTICIPANT DETAILS

#### Yeast strains and growth conditions

All experiments were performed in the prototrophic *Saccharomyces cerevisiae* CEN.PK background^43^. Gene deletions were generated by standard PCR-based disruption, in which drug-resistance markers bearing gene-specific flanking homology were integrated by homologous recombination. C-terminally tagged alleles were constructed analogously using PCR-amplified resistance cassettes with appropriate homology arms. Correct genome modifications were confirmed by DNA sequencing and by assessing the expected phenotypes. A complete list of strains and genetic manipulations is provided in the Key Resources Table. Cells were cultivated in synthetic (S) medium containing 0.67% yeast nitrogen base without amino acids (BD Difco), with or without 2% glucose as indicated. For solid media, yeast strains were grown on S agar supplemented with either 2% glucose (D), or 2% ethanol (E).

#### Nutrient switch experiments

For glucose depletion experiments, strains were pre-cultured overnight in SD medium. Overnight cultures were diluted into fresh SD to 0.2 OD/mL and allowed to grow for at least two doublings to reach logarithmic phase (OD600 ≈ 1). Acute glucose withdrawal was achieved by rapidly exchanging the medium: cultures were centrifuged, cell pellets were washed once with pre-warmed S medium, and cells were resuspended in an equal volume of pre-warmed S. Samples were harvested 30 mins after the shift. Unless otherwise noted, centrifugation steps were performed at 3000 rpm for 2 min at room temperature.

### METHOD DETAILS

#### Cell extracts preparation and western blot

Cells were precipitated by adding trichloroacetic acid to a final concentration of 10%, followed by an acetone wash of the pellets. Cell disruption was then carried out by bead beating in a denaturing urea buffer (6 M urea, 1% SDS, 50 mM Tris-HCl pH 7.5, 50 mM NaF, 5 mM EDTA) supplemented with 1 mM PMSF and a protease inhibitor cocktail (Roche). Clarified lysates were obtained by collecting the supernatant after heating the samples at 75 °C for 10 min. Protein amounts were quantified using the Pierce BCA assay, and equivalent protein quantities were resolved on NuPAGE gels. Membranes were blocked in 5% nonfat dry milk prior to antibody incubation.

#### RNA extraction and quantitative RT-PCR

RNA isolation of one OD600 unit of cells was performed using YeaStar RNA Kit with DNase treatment (Zymo Research) following the manufacturer’s protocol. A260 determined RNA concentration. 1 µg RNA was reverse transcribed to cDNA using Maxima H Minus cDNA Synthesis Master Mix with dsDNase from Thermo Fisher Scientific. Real-time PCR was performed in triplicate using SYBR Green Supermix from Bio-Rad. Transcript levels of genes were normalized to *TAF10A*. Primers are listed in Table S3.

#### Chromatin immunoprecipitation (ChIP)-sequencing

For ChIP assays, 100 OD600 units of cells were crosslinked in 1% formaldehyde for 15 min and quenched with 125 mM glycine for 10 min. Cell pellets were washed twice in buffer (100 mM NaCl, 10 mM Tris-HCl pH 8.0, 1 mM PMSF, 1 mM benzamidine-HCl) and snap-frozen in liquid nitrogen. Frozen cells were resuspended in 0.45 mL ChIP lysis buffer (50 mM HEPES-KOH pH 7.5, 500 mM NaCl, 1 mM EDTA, 1% Triton X-100, 0.1% deoxycholate, 0.1% SDS, 1 mM PMSF, 10 mM leupeptin, 5 mM pepstatin A, plus Roche protease inhibitor cocktail) and disrupted by bead beating. Lysates were divided into 100 µL portions and chromatin was sheared for 40 min in a PIXUL instrument (Active Motif) using the default settings (pulse number 50, frequency 1 kHz, burst rate 20 Hz). The sheared material was pooled and clarified by two spins at 15,000 rpm for 10 min each. For HA ChIP, chromatin was incubated overnight with 10 µg of anti-HA antibody. For spike-in normalization, exogenous chromatin and antibody were included following the manufacturer’s instructions (Active Motif). Immune complexes were captured with 25 µL magnetic beads for an additional 1.5 h. Beads were washed twice with ChIP lysis buffer, once with deoxycholate wash buffer (10 mM Tris-HCl, 0.25 M LiCl, 0.5% deoxycholate, 1 mM EDTA), and once with TE buffer (50 mM Tris-HCl pH 8.0, 10 mM EDTA). Bound chromatin was eluted in TES buffer (50 mM Tris-HCl pH 8.0, 10 mM EDTA, 1% SDS), and crosslinks were reversed at 65°C. Samples were then adjusted with an equal volume of TE containing 1.25 mg/mL proteinase K and 0.4 mg/mL RNase A and incubated for 2 h at 37°C before DNA purification with a QIAGEN PCR purification kit. Sequencing libraries were generated and 150 bp paired-end reads were obtained by Novogene.

### QUANTIFICATION AND STATISTICAL ANALYSIS

#### Analysis of ChIP-seq data

Analysis of raw data from all high-throughput sequencing experiments was performed using the Galaxy platform^44^. Sequencing adapters from the raw reads in FASTQ format were removed using Trim Galore using default settings. For spike-in normalization, reads were first mapped to the *Drosophila melanogaster* genome (dm6) using Bowtie2, and unmapped reads were then aligned to the *Saccharomyces cerevisiae* genome (sacCer3). The unmapped reads were removed using the tool Split BAM by reads mapping status. Counts per million (CPM) normalized bigWig tracks were generated using bamCompare while extending reads to the given average fragment size.

#### Peak calling

Peaks were called using MACS2 callpeak by normalizing against the pooled input and without building the shifting model. The pooled input file for +D and -D was generated by combining the data from 4 inputs each. Minimum FDR (q-value) cutoff for peak detection was 0.05. The noise from the data was reduced by using MACS2 bdgcmp. Next, peak refinement was performed with MACS2 bdgpeakcall using 5.0 as the cutoff, minimum length 200, and minimum gap 30 for peaks. Next, ChIPseeker was applied to obtain the genomic feature distribution under peaks, as well as the pie charts depicting that.

#### Downstream analysis

Integrative Genomics Viewer (IGV) was used for visualizing the genome browser tracks^45^. Functional profiling was performed using g:Profiler^46^. Venn diagrams were made using BioVenn^47^. Box plots and bar graphs were made using GraphPad Prism. Volcano plots and other graphs were made using Microsoft Excel.

#### Metagene Plots

The data was prepared for plotting using computeMatrix. As the output option either the scale-regions or the reference-point option was selected. For the scale region option, distance upstream and downstream of the region start/end position was selected as 2kb and 1kb, respectively. For the reference-point option, TSS was selected. Using the output of computeMatrix, plots were generated by plotHeatmap.

#### Analysis of RNA-seq data

Sequencing adapters from the raw reads in FASTQ format were removed using Trim Galore using default settings. Reads were aligned to the *Saccharomyces cerevisiae genome* (sacCer3) using HISAT2. For visualizing the read alignments, bigwig files were generated using the bamCoverage tool. For counting the reads per gene, featureCounts was used. DESeq2 was used to perform differential expression analysis.

---

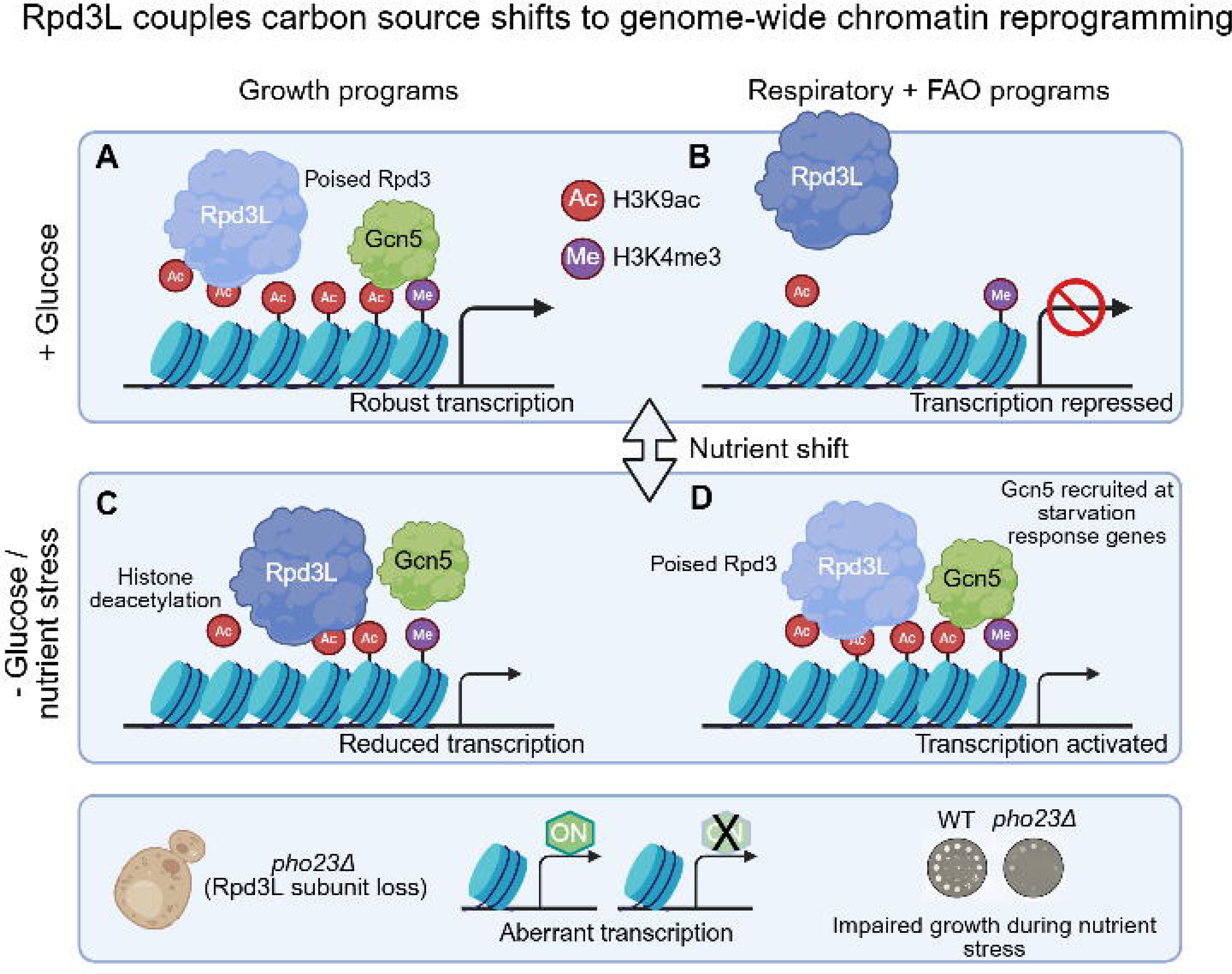

