## Supplementary figures for "Yeast Rpd3L histone deacetylase couples nutrient shifts to genome-wide chromatin reprogramming"

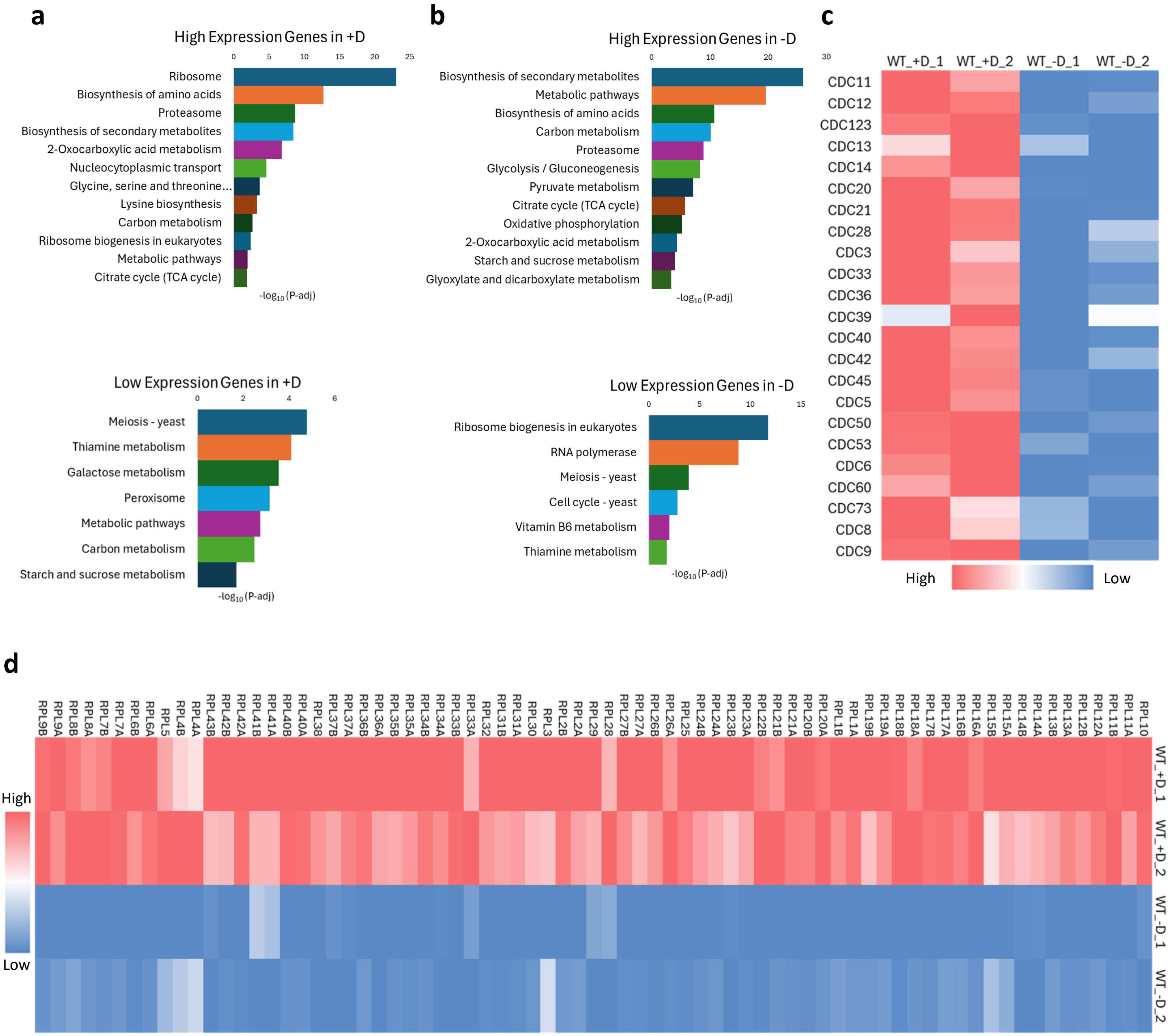


Supplementary figure S1: (a, b) KEGG analysis of highly or lowly expressed genes in glucose-replete (+D) or glucose-deplete (-D) media. (c, d) Heatmap showing the normalized counts of cell cycle (c) and ribosomal (d) genes from RNA-seq in +D and –D for the WT strain.

**a**

| Class | HDAC | Complex/Localization |
|---|---|---|
| Class I (Rpd3-like) | Rpd3 | Rpd3L, Rpd3S complexes |
| | Hos1 | Nuclear |
| | Hos2 | Set3C complex |
| | Hos3 | Cytoplasm, nuclear periphery |
| Class II (Hda1-like) | Hda1 | Hda1 complex (with Hda2/Hda3) |
| Class III (Sirtuin) | Sir2 | SIR complex (Sir2/3/4) |
| | Hst1 | Sum1/Rfm1/Hst1 complex |
| | Hst2 | Cytoplasmic & nuclear |
| | Hst3 | Nuclear |
| | Hst4 | Nuclear |

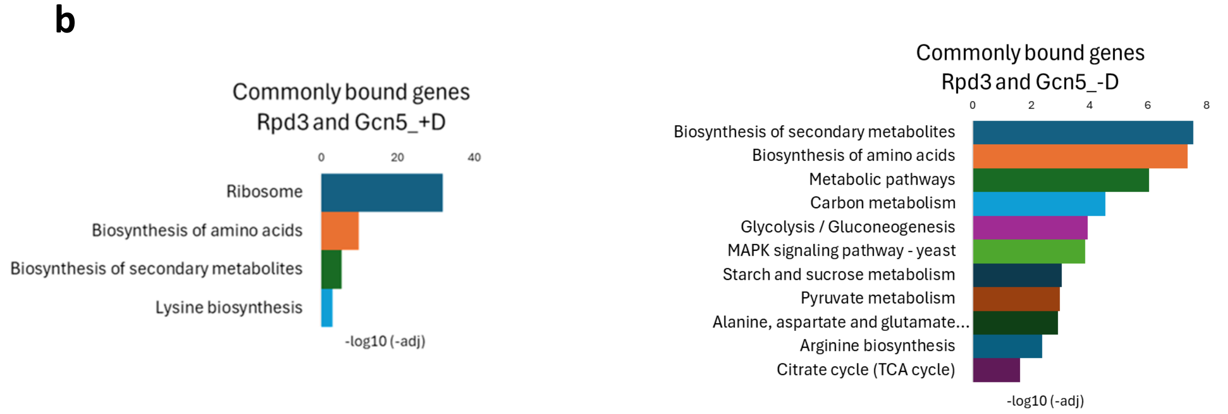


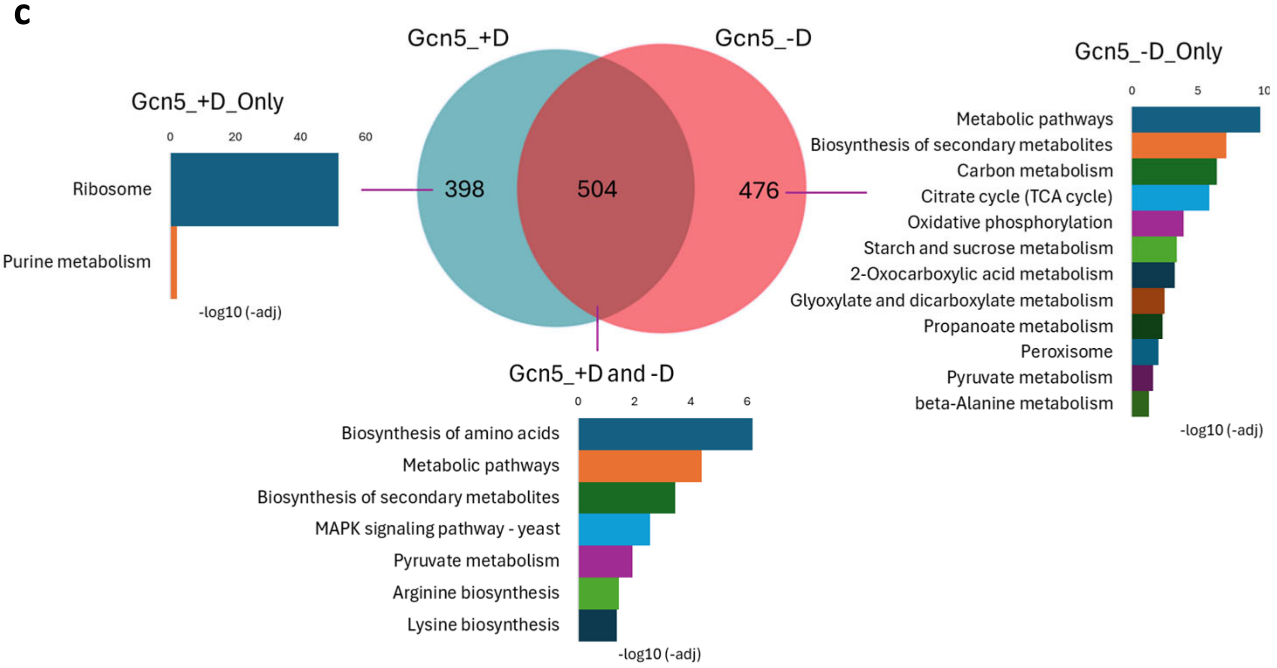


Supplementary figure S2: (a) HDACs in yeast. The HDACs highlighted in green were deleted from CEN.PK and the effect on H3K9ac level were tested by Western blotting. (b) KEGG analysis of commonly bound genes by Rpd3 and Gcn5 in +D and –D. (c) Venn diagram showing the overlap of genes that Gcn5 binds in +D vs -D. The three subsets were subjected to KEGG analysis, and the enriched pathways are shown.

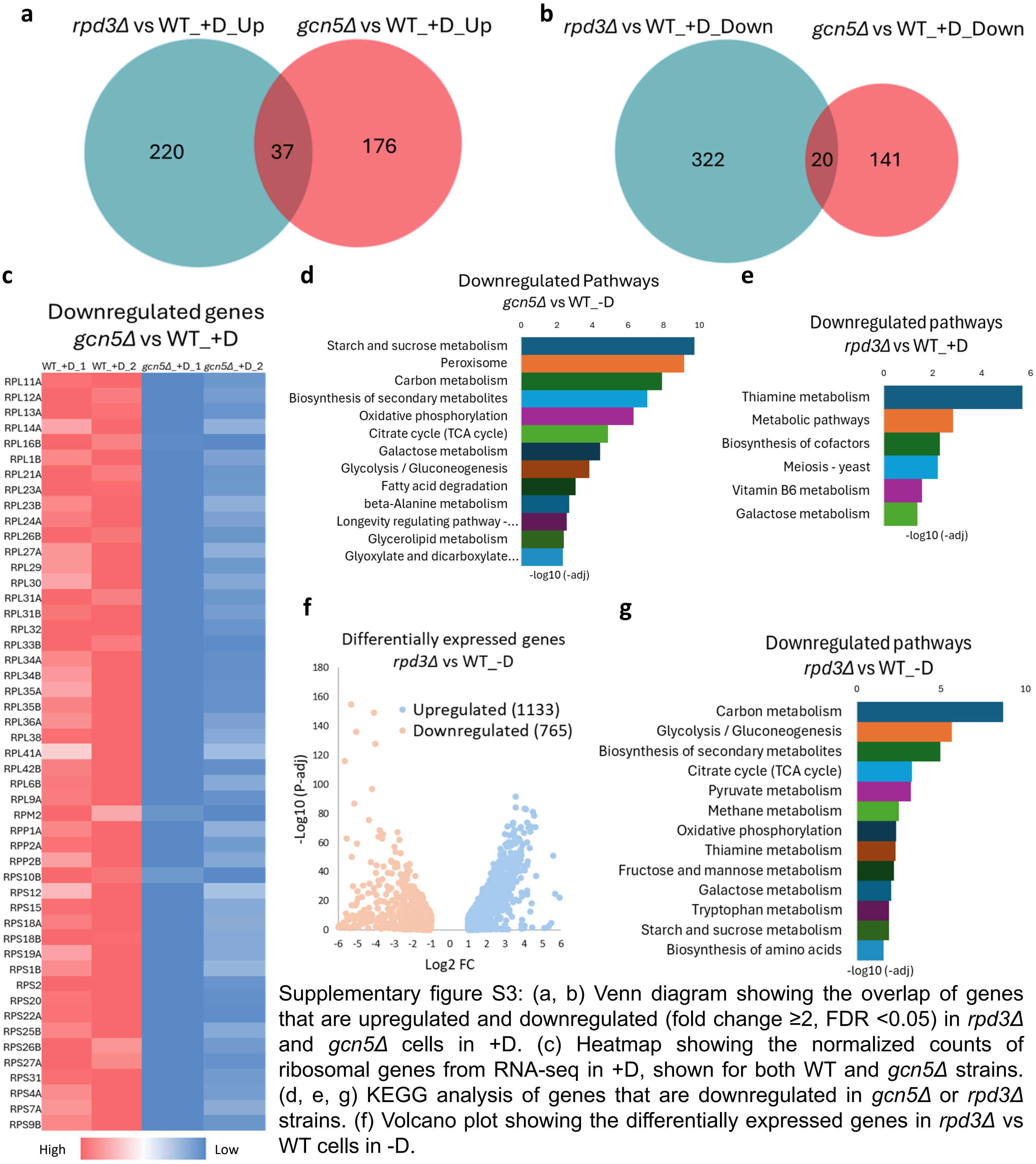


Supplementary figure S3: (a, b) Venn diagram showing the overlap of genes that are upregulated and downregulated (fold change ≥2, FDR <0.05) in *rpd3Δ* and *gcn5Δ* cells in +D. (c) Heatmap showing the normalized counts of ribosomal genes from RNA-seq in +D, shown for both WT and *gcn5Δ* strains. (d, e, g) KEGG analysis of genes that are downregulated in *gcn5Δ* or *rpd3Δ* strains. (f) Volcano plot showing the differentially expressed genes in *rpd3Δ* vs WT cells in -D.

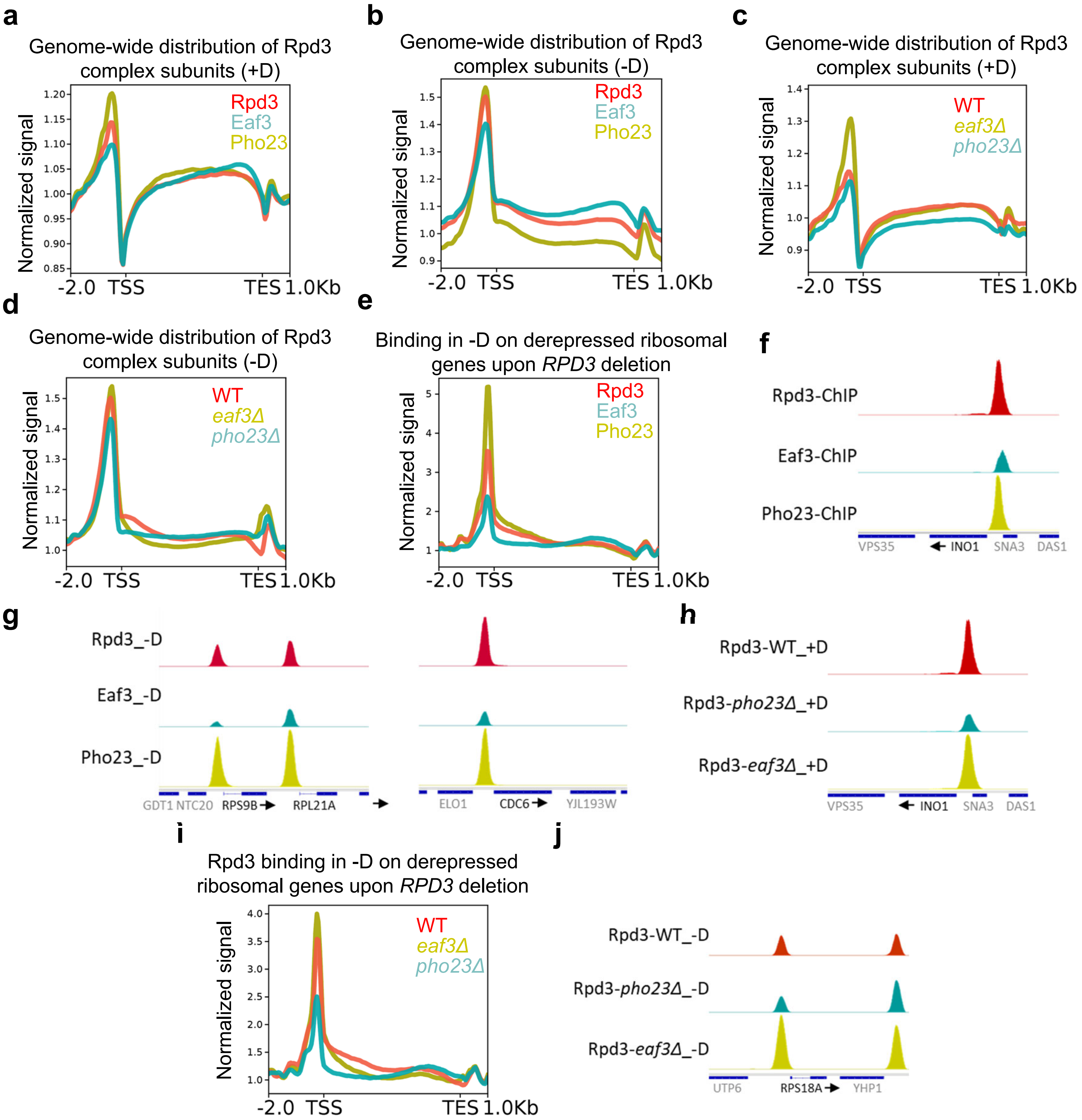


Supplementary figure S4: (a-e, i) Metagene profiles showing the genome-wide distribution of Rpd3, Eaf3, and Pho23 across all genes, as well as specifically at genes that are derepressed upon *RPD3* deletion in −D. (f-h, j) Representative genome browser tracks displaying ChIP-seq peaks for Rpd3, Eaf3, and Pho23 at selected loci.

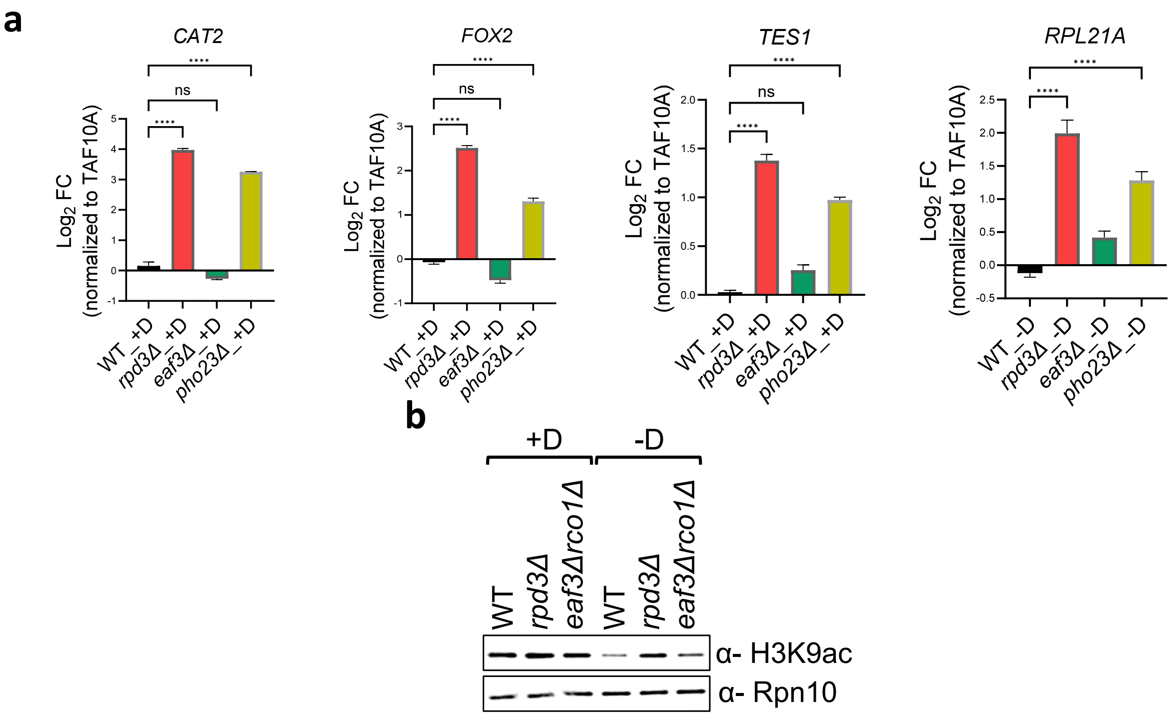


Supplementary figure S5: (a) qPCR analysis of representative genes in WT, *rpd3Δ*, *eaf3Δ*, and *pho23Δ*. Gene expression levels were normalized to an internal control, *TAF10A*. Ordinary one-way ANOVA was used to calculate statistical significance. n=3, ns>0.9999, ***p=0.0001, ****p<0.0001. The error bars depict the SEM. (b) Western blot showing H3K9ac levels in WT, *rpd3Δ*, and *eaf3Δrco1Δ* strain in +D and -D. Rpn10 was used as a loading control.

| Primer | Sequence 5' to 3' |
|---|---|
| CAT2_RT_F | GGCGGAAAAGCCGAACAAAT |
| CAT2_RT_R | TTCGGGCACGGGTAATGATG |
| POX1_RT_F | TGAGACAGACTTGCGGAGGA |
| POX1_RT_R | CTGAACCACCCAGTCGTCAT |
| RPL21A_RT_F | TTTCCAAACATTCGGTTATG |
| RPL21A_RT_R | ATCTGTTACCGACCATCTTG |
| TAF10_RT_F | ATATTCCAGGATCAGGTCTTCCGTAGC |
| TAF10_RT_R | GTAGTCTTCTCATTCTGTTGATGTTGTTGTTG |
| CRC1_RT_F | AGATGTCAAAATGGACAAGC |
| CRC1_RT_R | CCCTTAGAAGATGTTTGCAG |
| FOX2_RT_F | AGCATCCATATCAACTCTCG |
| FOX2_RT_R | GCGTTATTGACCAAGATGTC |
| RPS18A_RT_F | ACGCTTTGACCACTATCAAG |
| RPS18A_RT_R | TCTTCTACCAGTGGTCTTGG |
| TES1_RT_F | AACCTCGATGAGCAAGTATG |
| TES1_RT_R | GGTGAAAGTAAATCGTGTGG |

Table S2: qPCR primers
